# An evolutionarily conserved tryptophan cage promotes folding of the extended RNA recognition motif in the hnRNPR-like protein family

**DOI:** 10.1101/2024.12.20.629849

**Authors:** Ernest S. Atsrim, Catherine D. Eichhorn

## Abstract

The heterogeneous nuclear ribonucleoprotein (hnRNP) R-like family is a class of RNA binding proteins in the hnRNP superfamily that has diverse functions in RNA processing. Here, we present the 1.90 Å X-ray crystal structure and solution NMR studies of the first RNA recognition motif (RRM) of human hnRNPR. We find that this domain adopts an extended RRM (eRRM1) featuring a canonical RRM with a structured N-terminal extension (N_ext_) motif that docks against the RRM and extends the β-sheet surface. The adjoining loop positions the N_ext_ for docking to the RRM and forms a tryptophan cage motif, which has only been reported previously in a synthetic peptide. Using a combination of mutagenesis, solution NMR spectroscopy, and thermal denaturation studies we evaluate the importance of residues in the N_ext_-RRM interface and adjoining loop on eRRM folding and conformational dynamics. We find that these sites are essential for protein solubility, thermal stability, and conformational ordering. Consistent with their importance, mutations in the N_ext_-RRM interface and loop are associated with several cancers in a survey of somatic mutations in cancer studies. Sequence and structure comparison of the human hnRNPR eRRM1 to experimentally verified and predicted hnRNPR-like proteins reveal conserved features in the eRRM.

**Significance statement:** This study provides the first structural and thermodynamic analysis of the hnRNPR extended RNA recognition motif (eRRM1). Here an N-terminal extension docks alongside a canonical RRM, stabilized by an intervening loop containing a tryptophan cage motif. Highly conserved among hnRNPR-like proteins, this motif has only been previously observed in a synthetic peptide. Substitutions that disrupt observed interactions reduce thermal stability and are associated with cancers. These findings significantly advance our understanding of atypical RRM folding.

## Introduction

Heterogeneous nuclear ribonucleoproteins (hnRNPs) are RNA-binding proteins with diverse functions in cellular processes (1). Of these, the hnRNPR-like family is broadly involved in RNA processing and includes hnRNPR, hnRNPQ (also known as SYNCRIP and NASP1), dead end protein 1 (DND1), RBM46, RBM47, APOBEC1 complementation factor (A1CF), and GRY-RBP (2). hnRNPR plays important roles in splicing and transcription regulation (3–9). In mammals, the major isoform of hnRNPR consists of a predicted N-terminal α-helical bundle (NαB), three tandem RNA recognition motifs (RRMs), and a C-terminal RGG-box domain (10) (**Figure 1A**). A second isoform is also expressed in which the NαB is absent (11–13). Highly expressed in the nervous system (14), hnRNPR function is critical to axonal growth and neuronal development (15; 16; 5; 17). Deletion of RRM1 and RRM2 impaired β-actin content in growth cones and neurite growth (18).Overexpression and/or dysfunction of hnRNPR is linked to spinal muscular atrophy (19; 8; 20; 21) and cancer metastasis (22–24; 9). Mutations in the C-terminal RGG-box are associated with developmental disorders (25; 19). hnRNPR interacts with proteins such as Yb1 (13), SMN (14), HMGC (6), and ALS-associated proteins TDP-43 and FUS (20). hnRNPR has several identified RNA substrates including MHC class I mRNA (26), β actin mRNA (16), UPF3B mRNA (9), ASCL1 mRNA (21), and 7SK noncoding RNA (5; 27; 28; 7). Prior studies indicate that the first two RRMs are the primary RNA recognition sites (14; 18; 9).

**Figure 1.**
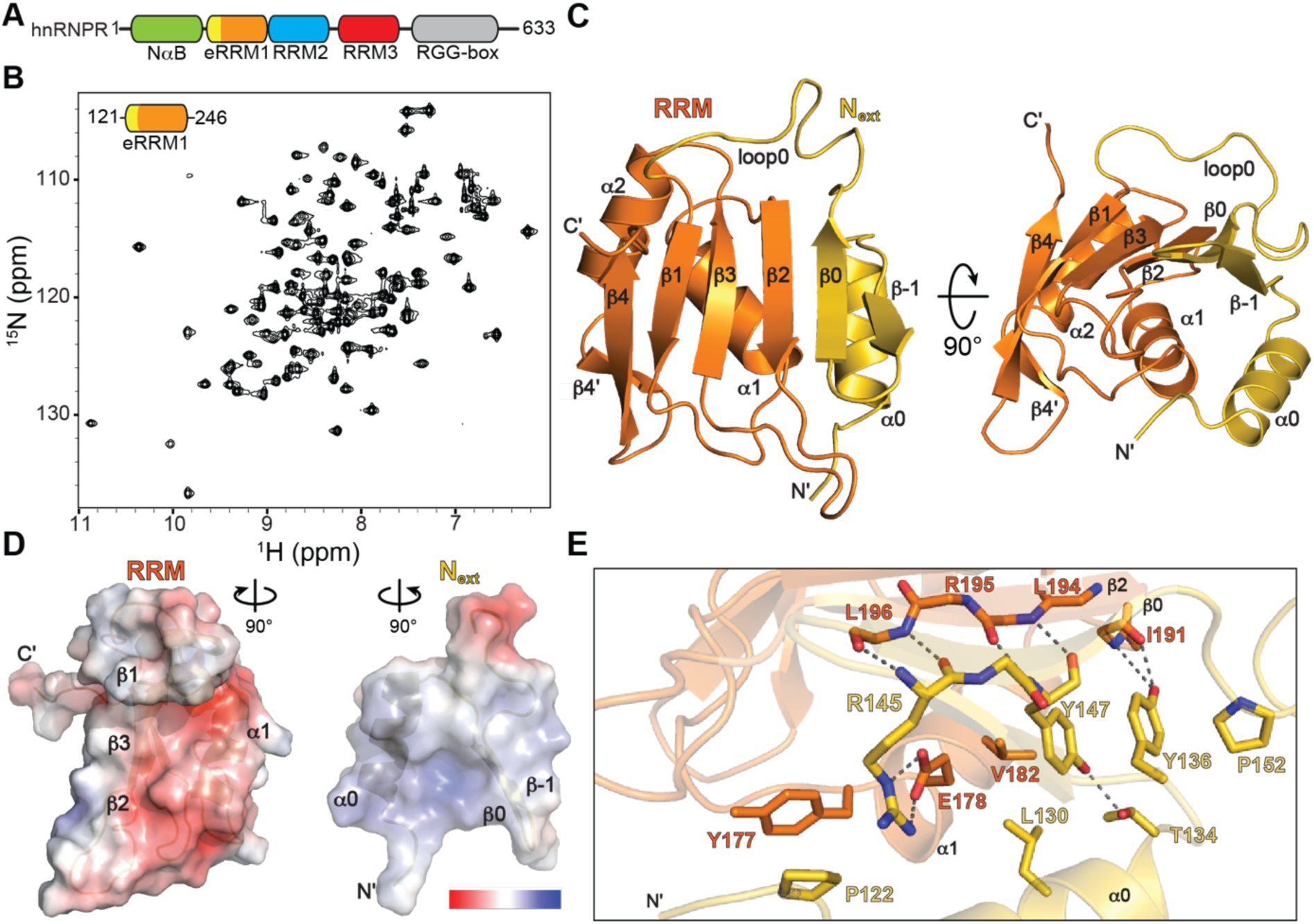
The first RNA binding domain of hnRNPR is an eRRM. A) Domain topology of human hnRNPR; B) ^1^H-^15^N HSQC spectrum of WT eRRM1 construct shows well-folded protein at 25 °C; C) 1.90 Å X-ray crystallographic structure of WT eRRM1 shows N_ext_ (colored gold) and RRM (colored orange) motifs; D) Electrostatic surface potential map of individual RRM and N_ext_ motifs show a charged interface; E) key residues participate in hydrogen bonding and hydrophobic interactions at the N_ext_-RRM interface.

Despite its significance, remarkably little is known regarding the structural and biophysical properties of hnRNPR. While an hnRNPR AlphaFold2 (AF2) predicted structural model is available, the only experimentally determined high-resolution structure is a solution NMR structural ensemble of RRM3 (PDB ID 2DK2). Canonical RRMs consist of an antiparallel β-sheet composed of four to five β-strands, with two α-helices that lie underneath the β-sheet (29). The β-sheet surface serves as the RNA recognition site, with conserved RNP sequence motifs on β1 (RNP2) and β3 (RNP1) that typically contain aromatic and/or basic residues (29). RRMs are frequently decorated with secondary structure elements at the N- and/or C-termini to promote RNA and/or protein substrate recognition and specificity (29–32). In particular, the U1-70K spliceosomal protein contains an RRM with an N-terminal α-helix, which improves RNA binding specificity and affinity (33). The first RRM of DND1 and hnRNPQ contain an N-terminal α-helix and β-hairpin, which contribute to the RNA binding surface (34; 35). All three atypical RRMs have been named extended RRMs (eRRMs) to denote their N-terminal secondary structure extension.

Here, we combine X-ray crystallography, solution NMR spectroscopy, thermal denaturation studies, and mutagenesis to determine the structure and conformational dynamics of the first RRM of the human hnRNPR (Hs-hnRNPR). We find that this domain is an eRRM consisting of an N-terminal extension (N_ext_) motif docked to a canonical RRM with structural similarity to hnRNPQ and DND1 eRRMs. Features include a structured loop connecting the N_ext_ to the RRM that organizes around a tryptophan residue in the β-sheet, forming a tryptophan cage motif that has only been previously reported in a synthetic peptide. Residues at the N_ext_-RRM interface as well as in the connecting loop are highly conserved among members of the hnRNPR-like family. N-terminal truncations or substitutions to residues in the N_ext_-RRM interface result in reduced protein solubility, reduced thermal stability, and increased conformational dynamics. Similarly, substitutions within the tryptophan cage result in reduced thermal stability and increased conformational dynamics, showing the importance of this motif in eRRM folding. Importantly, residues identified in this study to be important for protein folding are correlated with cancer-related missense mutations, suggesting a role in human health and disease. Together, this study characterizes features that define the eRRM1, identifies the tryptophan cage as a naturally occurring motif, and demonstrates the importance of stable association of the N_ext_ with the RRM for eRRM folding.

## Results

### Structure of the first RRM domain of hnRNPR is an extended RRM

Constructs of the first RRM domain of hnRNPR were designed with N- and C-terminal boundaries determined from sequence comparison to hnRNPR-like proteins and the AF2 predicted model (**Figure S1**). Human hnRNPR eRRM1 (WT eRRM1, residues 121-246) containing an N-terminal histidine tag and TEV cleavage site was overexpressed in *E. coli* and purified (see methods). The solution state NMR ^1^H-^15^N heteronuclear single quantum coherence (HSQC) spectrum showed excellent chemical shift dispersion indicating a stably folded protein (**Figures 1B and S2**). NMR resonance assignments were performed using standard triple-resonance experiments, with 94.83% completeness for backbone resonances (N, H, Cα, Cβ, CO). Residues D179-L181 (α1) could not be assigned due to line broadening. We performed crystal screening of constructs for X-ray crystallographic structure determination and identified conditions yielding crystals that diffracted to 1.90 Å (**Table S1**).

WT eRRM1 folds as an eRRM containing a canonical RRM domain with an N-terminal extension (**Figure 1C**). The canonical RRM contains a β_1_α_1_β_2_β_3_α_2_β_4’_β_4_ topology and is comprised of a β-sheet with five anti-parallel β-strands and two α-helices that lie underneath the β-sheet. This secondary structure is consistent with predictions made from chemical shift assignments using TALOS+ (36; 37) (**Figure S3**). On the β-sheet surface, the RRM domain has conserved RNA recognition sequences RNP2 on β1 (V167-F168-V169) and RNP1 on β3 (Y208-A209-F210) (38). An 18 aa structured loop (loop0, G149-T165) links β0 in the N_ext_ to β1 in the RRM. The N_ext_ contains an α_0_β_-1_β_0_ topology and is comprised of an α-helix that lies underneath a β-hairpin. The N_ext_ β-hairpin docks alongside the β2-α1 side of the canonical RRM, extending the β-sheet surface by two β-strands. The N_ext_-RRM interface is highly charged, with an electronegative surface potential on the β2-α1 side of the RRM and an electropositive surface potential on the α0-β0 side of the N_ext_ (**Figure 1D**). There are several interactions between N_ext_ and RRM subunits including hydrogen bonds between residues G143 (loopβ-1-β0)-M198 (β2), Q144 (β0)-R195 (β2), R145 (β0)-L196 (β2), R145 (β0)-E178 (α1), Y147 (β0)-L198 (β2), and G149 (loop0)-D193 (β2) as well as hydrophobic contacts between residues P122-Y177, L130-V182, Y136-P152, and Y136-I191 (**Figure 1E**).

### A structurally conserved tryptophan cage stabilizes loop0 connecting N_ext_ to RRM

In the X-ray crystal structure, the 18 aa loop0 that links the N_ext_ to the canonical RRM is highly structured, where primarily hydrophobic residues are coordinated around β_2_ residue W192 with extensive van der Waal (vdW) contacts (**Figure 2A**). The W192 indole ring is positioned between P150-P151 and Q160-P161, has an edge-to-face interaction with Y156, and contacts D193 (β2) and T212 (β3) on the β-sheet (**Figure 2A**). This organization bears a striking resemblance to the tryptophan cage (Trp-cage), a synthetic peptide derived from the exendin-4 peptide (39) (**Figure 2B**). The Trp-cage also has a central tryptophan residue (W6) positioned between two proline residues and has a similar edge-to-face interaction with a tyrosine residue (Y3). Rather than residing on a β-sheet as observed in eRRM1, W6 is in an α-helix and interacts with α-helical residues Y3 and L7. However, the positioning of Y3 and L7 sidechains are nearly identical to T212 and D193 on eRRM1, respectively, despite the significant secondary structure differences. An X-ray crystal structure reported for the *Drosophila melanogaster* hnRNPQ (Dm-hnRNPQ) eRRM1 domain has near-identical loop0 sequence and structural similarity to Hs-hnRNPR (**Figure 2C**) (34). Here, residue W155 in Dm-hnRNPQ replaces residue Y156 in Hs-hnRNPR for maintained edge-to-face interactions with the β2 residue W191 in Dm-hnRNPQ. A valine (V159) is positioned above W191 rather than Q160 in Hs-hnRNPR for maintained vdW contacts with W191 (**Figure 2C**).

**Figure 2.**
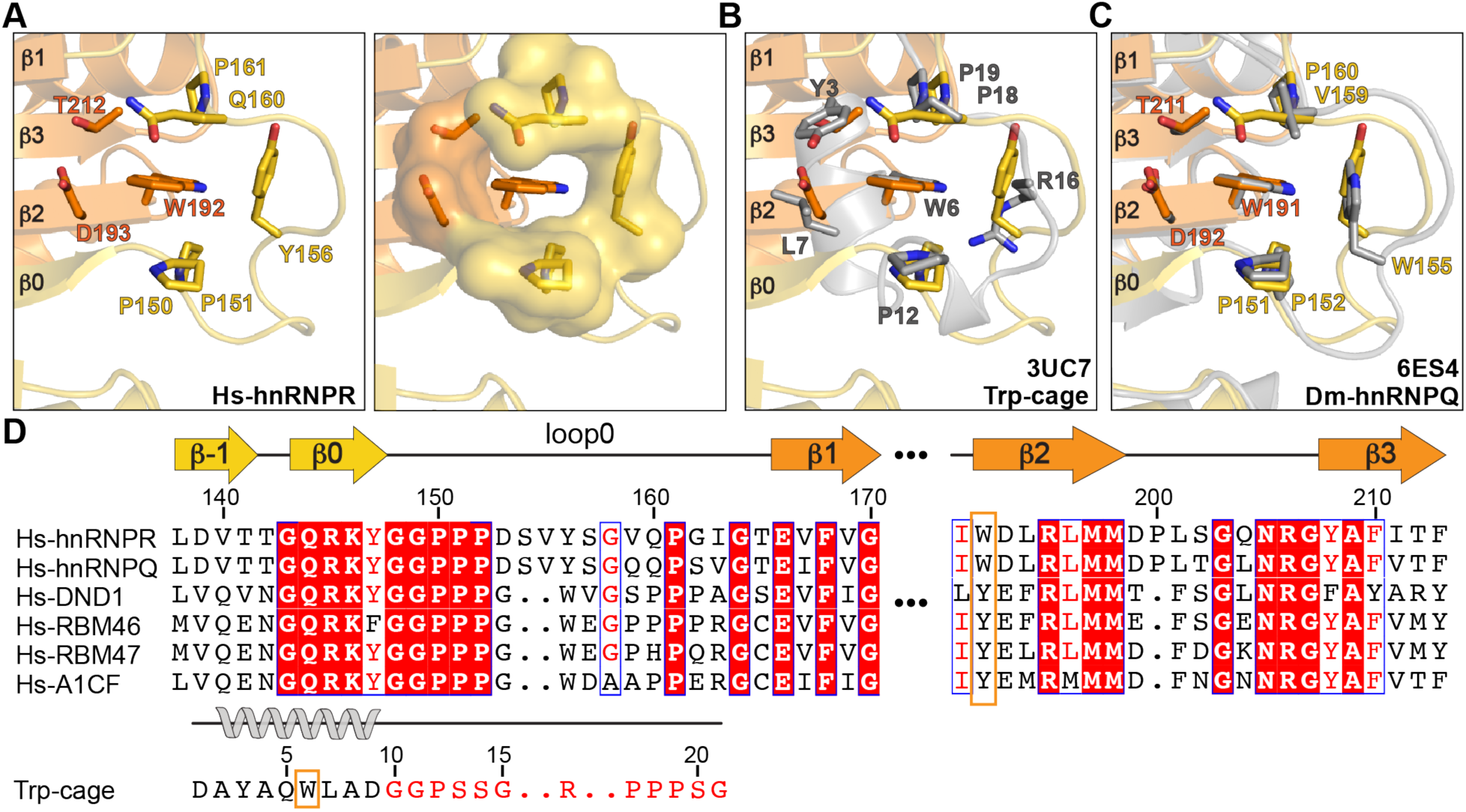
A conserved tryptophan cage motif structures the N_ext_ loop0. A) hnRNPR eRRM1 residues in loop0 and the β-sheet stabilize loop0 *left*: sidechains encircling W192 are shown in surface representation; B) Hs-hnRNPR eRRM1 superimposed with synthetic Trp-cage, colored gray (PDB ID 3UC7, chain C) (63), C) Hs-hnRNPR eRRM1 superimposed with Dm-hnRNPQ eRRM1, colored gray (PDB ID 6ES4, chain A) (34); D) Sequence alignment of eRRM1 domains among human hnRNPR-like proteins shows high conservation for residues at the N_ext_-RRM interface and tryptophan cage. Trp-cage secondary structure and sequence are shown below the hnRNPR-like family sequence alignment and show high sequence conservation in the loop. The central tryptophan residue is indicated in an orange outline.

To gain insights into the sequence conservation of the tryptophan cage motif across the hnRNPR-like family of proteins, we compared the synthetic Trp-cage sequence to human hnRNPR, hnRNPQ, DND1, RBM46, RBM47, and APOBEC1 complementation factor (A1CF) proteins (**Figure 2D**). In all cases, GGP is found at the transition from the secondary structured element to the loop, promoting a sharp turn in loop0. In addition, a proline is conserved at position 161, positioned above the tryptophan residue. In humans, hnRNPR and hnRNPQ contain a two aa insertion in loop0 compared to DND1, RBM46, RBM47, and A1CF. Notably, residues in loop0 are conserved including an aromatic residue at position 156. While a glutamine is conserved in hnRNPR and hnRNPQ at position 160, a proline is located at this position in the synthetic Trp-cage and other members of the hnRNPR-like family. Interestingly, DND1 and RBM46 both contain a triple proline repeat sequence at residues 160-162, in common with the synthetic Trp-cage at residues 17-19.

To identify sequence conservation across hnRNPR proteins, we curated a list of ten known and 215 predicted/hypothetical hnRNPR proteins identified in a Uniprot BLAST search (see methods). Among the ten known hnRNPR sequences, eRRM1 contains near-exact sequence identity (**Figure S4**). In particular, residues at the N_ext_-RRM interface and tryptophan cage in loop0 are identical. Sequence similarity is conserved at positions 154, 215, and 228 as polar; position 159 as hydrophobic; and position 217 as negatively charged. The only dissimilarity occurs at position 187, which is conserved as a lysine across all species except in *Rattus norvegicus*, where it is a glutamic acid. We expanded our sequence analysis to include predicted/hypothetical hnRNPR sequences and observed similarly high conservation in the eRRM1, particularly in the N_ext_ and N_ext_-RRM interface (**Figure S5**). Within loop0, the N- and C-terminal ends are highly conserved while central residues at position 154-157 are variable. In addition, the loop between β1 and α1 is highly variable. Taken together, both the sequence and structure of loop0 are highly conserved as a tryptophan cage motif in the eRRM1. In particular, the tryptophan residue in β2 has a dual role where the backbone participates in N_ext_-RRM interface interactions and the indole sidechain participates in the tryptophan cage motif. To our knowledge, this study reports the first example of a tryptophan cage motif naturally occurring in a biologically relevant protein.

### Conserved residues at the N_ext_-RRM interface and loop0 are required for protein folding stability

We next investigated the impact of truncations and point substitutions on eRRM1 chemical environment and thermal stability, evaluated using NMR chemical shift perturbation (CSP) analysis and thermal denaturation studies. The WT eRRM1 construct has a melting temperature (Tm) of 59 ± 1 °C and a highly cooperative melting transition (**Figures 3A and S2B, Table 1**). To determine the importance of the N_ext_ motif in the eRRM1, we generated a construct of the canonical RRM domain lacking N_ext_ (aa 162-246). This construct was primarily observed in the insoluble fraction in *E. coli,* with little soluble expression (**Figure S6**). We next extended the N-terminus to residue 116 (Ext), which includes the full linker region between the NαB and eRRM1 domains. In the backbone amide ^1^H-^15^N HSQC spectrum, the spectra for Ext and WT eRRM1 constructs were nearly identical (**Figure S7**). Four additional resonances were observed for residues E117-K120 that could be assigned from the ^15^N-edited NOESY spectrum. CSPs were observed for residues G121-D123 (N-terminus), D175-L176 (loopβ1-α1), and G203-N205 (loopβ2-β3) (**Figure S7A-C**), consistent with interactions observed in the X-ray crystal structure. This construct showed a modest increase in Tm compared to WT (ΔTm 3.0 ± 1.4 °C) (**Figures 3A and S7D**). Together, these data are consistent with the N-terminal boundary of the eRRM1 beginning at residue 121.

**Figure 3.**
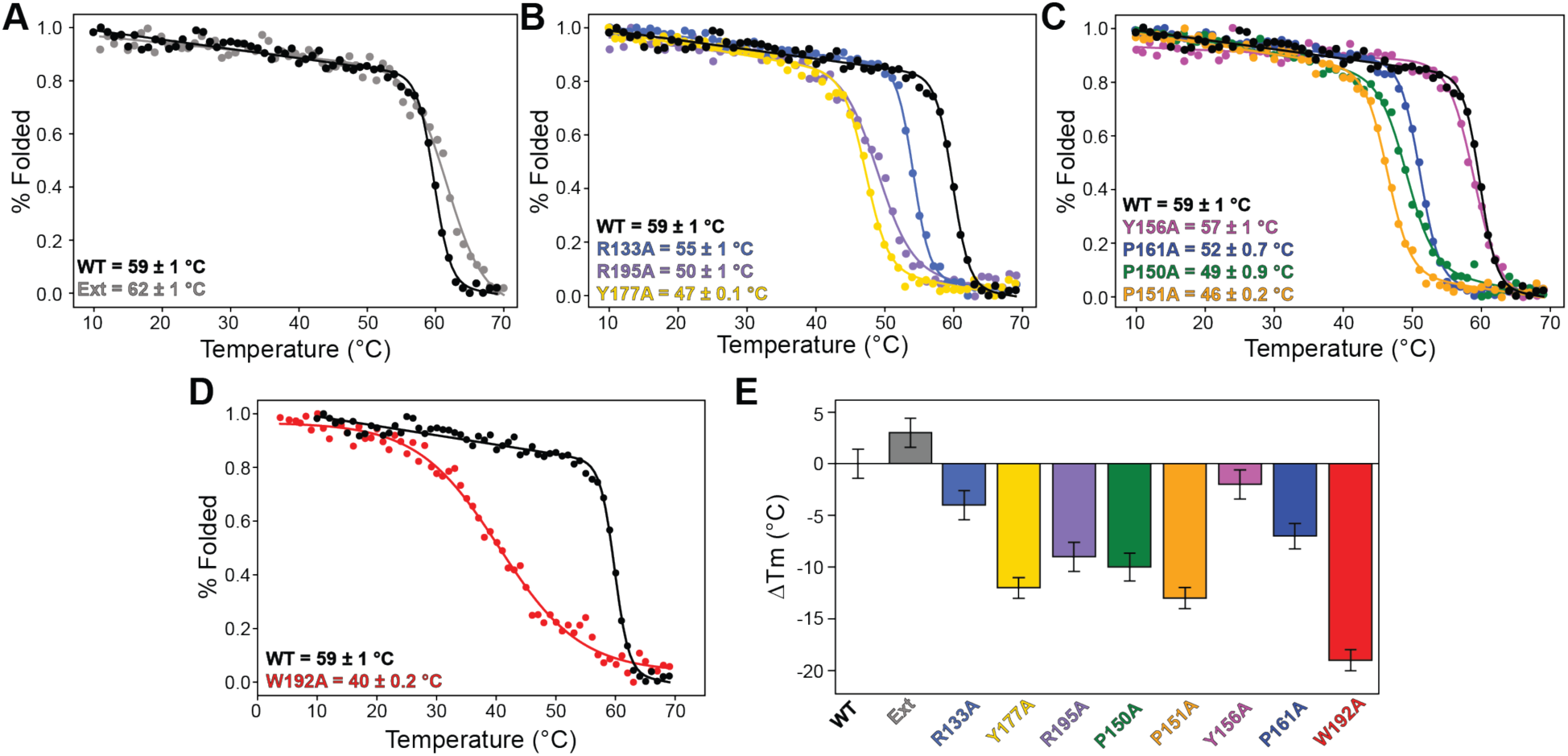
Impact of sequence variation on eRRM1 thermal stability using thermal denaturation studies. A-C) Thermal denaturation plots for A) WT (colored in black) and Ext (colored in gray) constructs; B) alanine substitutions to residues at N_ext_-RRM interface; C) alanine substitution of loop0 residues in the tryptophan cage motif; D) W192A construct (colored in red) shows severe reduction in thermal stability; E) Bar plot summarizing melting temperature data for constructs compared to WT eRRM1.

**Table 1.** Thermal melting statistics.

| **Construct** | **Protein boundary (aa)** | **T<sub>m</sub> (°C)** |
| --- | --- | --- |
| WT | 121-246 | 59 ± 1 |
| **N-terminal extension and truncation** |  |  |
| Ext | 116-246 | 62 ± 1 |
| RRM1 | 162-246 | ND |
| **Point substitutions at the N<sub>ext</sub>-RRM1 interface** |  |  |
| R133A | 121-246 | 55 ± 1 |
| R145A | 121-246 | ND |
| R195A | 121-246 | 50 ± 1 |
| Y177A | 121-246 | 47 ± 0.1 |
| **Point substitutions in loop0** |  |  |
| P150A | 121-246 | 49 ± 0.9 |
| P151A | 121-246 | 46 ± 0.2 |
| Y156A | 121-246 | 57 ± 1 |
| P161A | 121-246 | 52 ± 0.7 |
| W192A | 121-246 | 40 ± 0.2 |

*ND due to insoluble expression*

In order to evaluate the contribution of specific residues at the N_ext_-RRM interface on protein thermal stability, we next performed alanine substitutions to residues R133 (α0), R145 (β0), Y177 (loopβ1-α1), and R195 (β2) (**Figure 3B and Table 1**). R133 (α0) is positioned toward helix α1 but does not make direct contacts to the canonical RRM domain in the X-ray crystal structure. Alanine substitution (R133A) shows significant CSPs for N_ext_ α-helical residues E124, R133A, and T134, and to a lesser extent RRM residues F185-K187 (α1), G189 and I191 (loopα1-β2), and W192 (β2) (**Figure S8A-C**). A modest reduction in the Tm (ΔTm −4.0 ± 1.4 °C) was observed (**Figure 3B and S8D**). The R145 (β0) sidechain has extensive interactions that span both N_ext_ and RRM domains including a cation-π interaction with Y177 (loopβ1-α1) and salt bridge to E178 (α1) (**Figure 1E**). Consistent with its unique environment, the Hε resonance is significantly downfield shifted (10.8 ppm) relative to other arginine Hε resonances in the ^1^H-^15^N HSQC spectrum, indicating decreased shielding (**Figure S9**). R145A does not express in *E. coli*, indicating the importance of these interactions on protein solubility. Alanine substitution of Y177 (Y177A) showed significant CSPs for L176-Y177A, at the substitution site, and to a lesser extent residues E124-Y147 (α0) (**Figure S10A-C**). Y177A has a significantly reduced Tm (ΔTm −12 ± 1.0 °C) (**Figure 3B and S10D**). The R195 (β2) sidechain is positioned on the β-sheet surface and hydrogen bonds to Q144 (β0), D193 (β2), T212 (β3), E166 (β1), and a coordinated water molecule (**Figure S11**). Consistent with these interactions, R195A showed CSPs across the β-sheet surface, particularly for residues G143-R145 (β0), Y147 (α0), V167-F168 (β1), and D193-L196 (β2) near the substitution site (**Figure S11A-C**). R195A has a significantly reduced Tm (ΔTm −9.0 ± 1.4 °C) compared to WT (**Figure 3B and S11E**). Together, these data support the importance of stable association between N_ext_ and RRM subunits for protein solubility and thermal stability.

We next performed alanine substitutions to residues in loop0 to investigate the importance of conserved residues in the tryptophan cage on eRRM1 thermal stability (**Figure 3C**). Substituting Y156 to alanine (Y156A) resulted in significant CSPs for residues D153-S154 (loop0), G162-I163 (loop0), and C214 (loopβ3-α2) (**Figure S12A-C**). A minor reduction was observed in Tm relative to WT (ΔTm −2.0 ± 1.4 °C) (**Figure 3C and S12D**). P161A shows significant CSPs for residues I163-T165 (loop0), T212 (β3), and G215 (β3) (**Figure S13A-C**), with a modest reduction on Tm (ΔTm −7.0 ± 1.2 °C) (**Figures 3C and S13D**). In the P150A construct, CSPs were observed for residues T137-L138 (β-1), Y147-G149 (loop0), and S157 (loop0) (**Figure S14A-C**). Similar CSPs were observed for the P151A construct, particularly residues G149, Y156, G158, Q160 (loop0) (**Figure S15A-C**). In the P151A construct, residues W192 and L194 also showed significant CSPs. Both P150A and P151A constructs show substantial reductions of Tm (ΔTm - 10.0 ± 1.3 °C and ΔTm −13 ± 1.0 °C, respectively) (**Figures 3C, S14D, S15D**).

W192 is located both at the N_ext_-RRM interface, with backbone interactions to G149 (β0), and in the tryptophan cage as the central residue. Alanine substitution of the central W192 (β2) residue in the tryptophan cage (W192A) dramatically reduced sample stability, resulting in precipitation within 24 hours and a severe reduction in thermal stability (ΔTm −19 ± 1.0 °C) with reduced cooperativity (**Figures 3D and S16**). Significant CSPs are observed in loop0 and across the β-sheet, particularly in β3 and β1 strands (**Figure S16A-C**). In summary, all point substitutions that disrupt observed sidechain interactions at the N_ext_-RRM interface and loop0 result in reduced thermal stability (**Figure 3E**). Y177A, P151A, and W192A substitutions showed the greatest reductions in thermal stability, indicating their significance in eRRM1 folding through stabilizing the N_ext_-RRM interface and tryptophan cage motif interactions, respectively. Together, these results show the importance of stable formation of the N_ext_-RRM interface and tryptophan cage motif on eRRM1 protein folding.

### Point substitutions enhance conformational dynamics in loop0 and N_ext_-RRM interface

We used solution NMR spectroscopy to investigate WT eRRM1 conformational dynamics at fast (ps-ns) timescales. Longitudinal (R_1_), and transverse (R_2_) relaxation rates were measured for amide nitrogen resonances, and ^1^H-^15^N heteronuclear NOE values for backbone amide resonances, at 25 °C (**Figure 4A-C**). These data together show an overall highly ordered protein with a flexible N-terminal residue G121 and C-terminal residues N245-N246. Similarly, ^1^H-^15^N NOE values of the Ext construct gradually reduce approaching the N-terminus, with a minimum value for residue E117, consistent with G121 as the N-terminal boundary of the eRRM1 (**Figure S7E**). ^1^H-^15^N NOE values are slightly elevated for residues K146-G148 (β0) in Ext compared to WT, indicating that the additional N-terminal residues result in increased order in β0 at the N_ext_-RRM interface. N-terminal residue D123 (α1) has similar values as structured regions, consistent with the hydrophobic contact observed between P122 and Y177 (α1) in the X-ray crystal structure (**Figure 1E**). Loop residues V159-G164 (loop0) and L201-Q204 (loopβ2-β3) have reduced values, indicating these loops have fast internal motions. Elevated R_2_ values are observed for residues Y147 (β0) and V167 (β1), located at β-strand ends that connect to loop0, as well as residue W192 (β2), located in the tryptophan cage. These elevated values indicate additional contributions to R_2_ from chemical exchange, which may be due to slow motions in the β-strand residues connected to loop0 and at the N_ext_-RRM interface.

**Figure 4.**
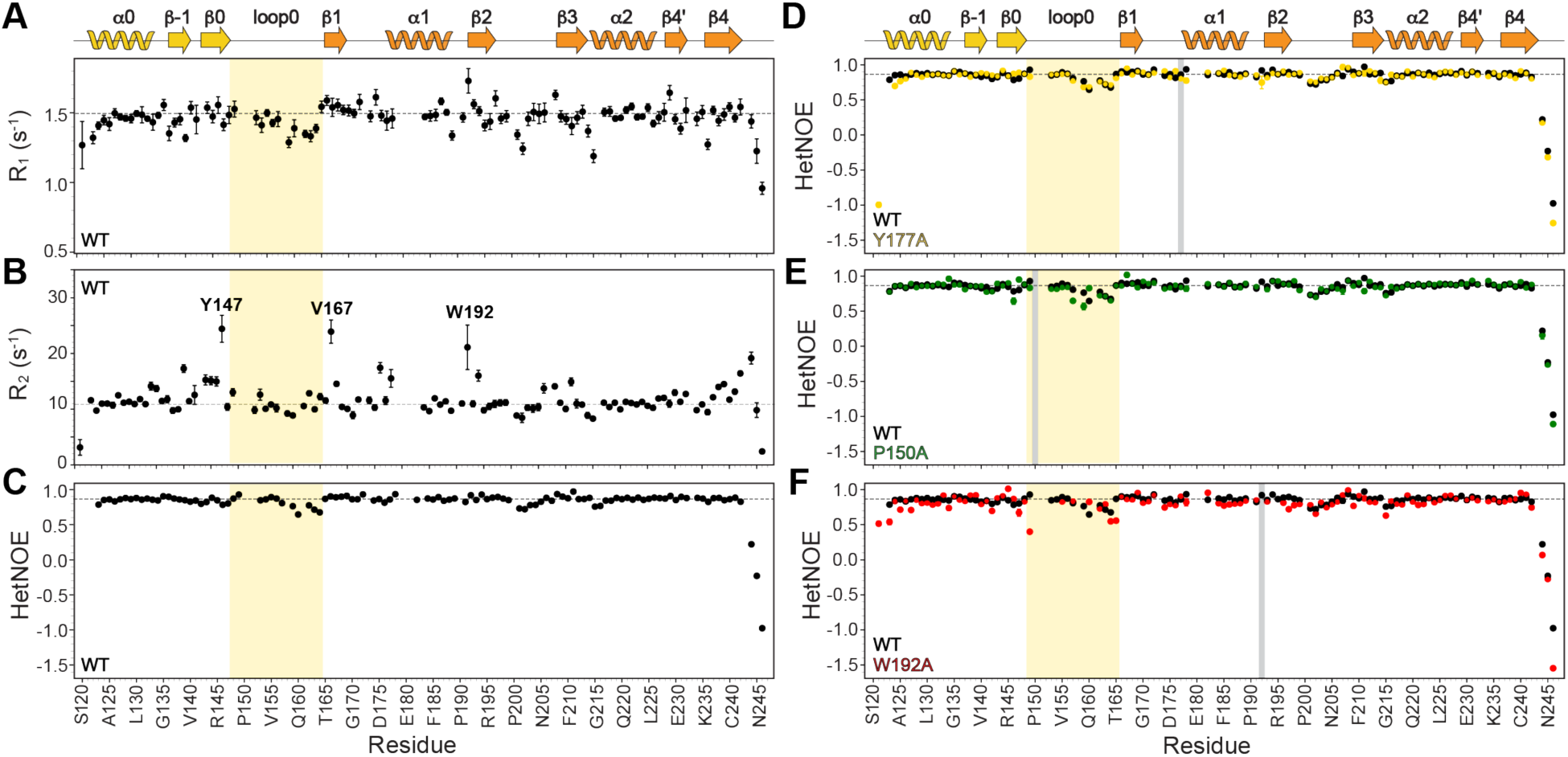
hnRNPR eRRM1 conformational dynamics show an ordered protein with flexible ends. A-C) Plots of WT eRRM1 A) R_1_ relaxation rates; B) R_2_ relaxation rates, with labeled residues indicating sites of chemical exchange; C) ^1^H-^15^N NOE values. D-F) ^1^H-^15^N NOE values for WT eRRM1 (colored in black) and point substitution constructs D) Y177A (colored in yellow), E) P150A (colored in green), and F) W192A (colored in red). The eRRM1 secondary structure topology is depicted above panels A and D. The substitution site is shown as a gray bar, and loop0 is highlighted in light yellow.

We next compared the conformational dynamics of WT eRRM1 to the point substitution constructs described above to evaluate the impact on eRRM1 conformational dynamics. In the N_ext_-RRM interface, ^1^H-^15^N NOE values of R133A and R195A constructs are nearly identical to WT, indicating no change in fast internal motions (**Figures S8E and S11F**) despite both constructs having reduced thermal stability compared to WT (**Figure 3E**). In contrast, in the Y177A construct ^1^H-^15^N NOE values are reduced for N-terminal residues E124-A125 (α0) relative to WT indicating increased mobility at the N-terminus, particularly in helix α0 (**Figure 4D**). These increased dynamics are likely due to destabilizing the hydrophobic contact between P122-Y177A and are consistent with the substantial reduction in thermal stability compared to WT.

We next compared ^1^H-^15^N NOE values for loop0 point substitution constructs to evaluate their impact on eRRM1 conformational dynamics. ^1^H-^15^N NOE values in the Y156A construct are nearly identical to WT, indicating no change in relative internal motions and consistent with this construct’s similar thermal stability to WT (**Figures 3E and S12E**). In the P161A construct, ^1^H-^15^N NOE values are slightly reduced for loop0 residues V159-Q160 and G162 that are adjacent to the substitution site (**Figure S13E**). Similarly, in both P150A and P151A constructs ^1^H-^15^N NOE values are reduced for loop0 residues S157-Q160 (**Figures 4E and S15E**), indicating increased dynamics compared to WT. The W192A construct, which has the most deleterious impact on thermal stability (**Figure 3E**), showed globally increased dynamics compared to WT (**Figure 4F**). In particular, ^1^H-^15^N NOE values are reduced for residues at the N-terminus, loop0, and β2 indicating increased dynamics at the N_ext_-RRM interface and loop0 as a result of removing the central tryptophan indole ring. These results indicate that the reduced thermal stability observed in point substitution constructs is due to increased dynamics in loop0 and reduced association between N_ext_ and RRM motifs.

### Missense mutations in N_ext_ motif are associated with cancer

Prior literature has shown disease-associated mutations in hnRNPR at RRM2 (9) and the C-terminal RGG-box repeat (25; 19). To investigate the biomedical relevance of eRRM1, we surveyed the COSMIC (Catalog of Somatic Mutations in Cancer) database to map sequence variants onto hnRNPR domains (40; 41). Of the 513 variant sequences present, there were 172 missense, or single amino acid residue, mutations (**Supplemental File 2**). Missense mutations were present in all domains, with 38.4% in the RGG-box, 16.9% in eRRM1, 16.3% in RRM3, 12.8% in RRM2, 12.2% in NαB, and 3.4% in linker regions (**Figure S17A-B**). Missense mutations are associated with numerous cancer histology subtypes, most frequently adenocarcinoma (38.3%), not specified (28.8%), and squamous cell carcinoma (11.9%) (**Figure S17C**). Within the eRRM1 domain, missense mutations are most frequently present in the adenocarcinoma cancer subtype (37.3%) and are found in the large intestine (21.6%) and skin (15.7%) (**Figure S17D-E**). Of the 27 unique missense mutations in eRRM1, 13 were present in the N_ext_ motif, loop0, and N_ext-_RRM1 interface residues including at residues P122, T134, G135, G149, P152, Y156-G158, T165-E166, and R195 (**Figure 5**). Our sequence conservation analysis and mutagenesis studies showed that several of these residues are highly conserved and required for protein thermal stability. Together, these data suggest that destabilization of the N_ext-_RRM interface is associated with cancer.

**Figure 5.**
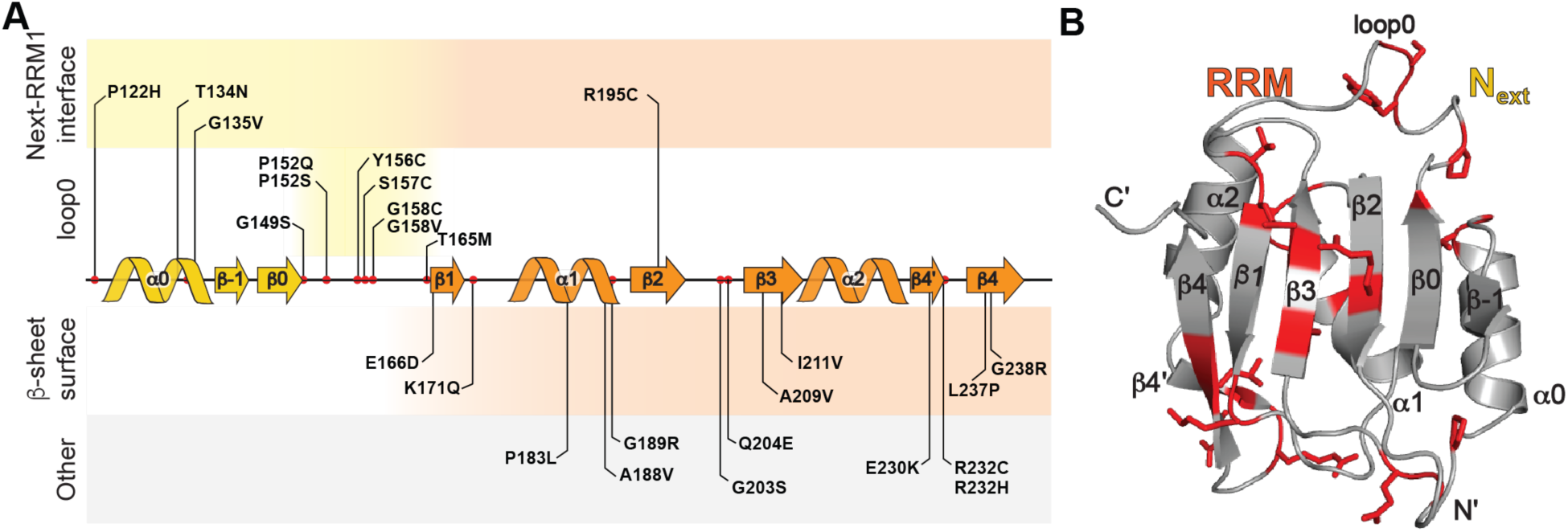
Missense mutations in the N_ext_-RRM interface and loop0 are associated with cancers. A) COSMIC-annotated missense mutations mapped onto the eRRM1 secondary structure topology, B) COSMIC-annotated missense mutations (colored in red with sidechains shown in stick representation) mapped onto the X-ray crystal structure of the hnRNPR eRRM1 domain.

### Comparison of human hnRNPR eRRM1 to hnRNPR-like protein family

The Hs-hnRNPR eRRM1 X-ray crystal structure is extremely similar to the predicted AF2 model, with an overall backbone RMSD of 0.31 Å (**Figure 6A**). Comparison to the previously determined X-ray crystal structure of the Dm-hnRNPQ eRRM1 (34) also shows high similarity, with an overall backbone RMSD of 0.50 Å (aa 122-242) (**Figure 6B**). Differences are observed in loop0, helix α0 orientation, and positioning of the N_ext_ β-turn (**Figure 6B**), which may be due to a 2 aa deletion between the triple proline repeat and aromatic residue in the eRRM1 of Dm-hnRNPQ compared to human hnRNPR and hnRNPQ (**Figure 2D**).

**Figure 6.**
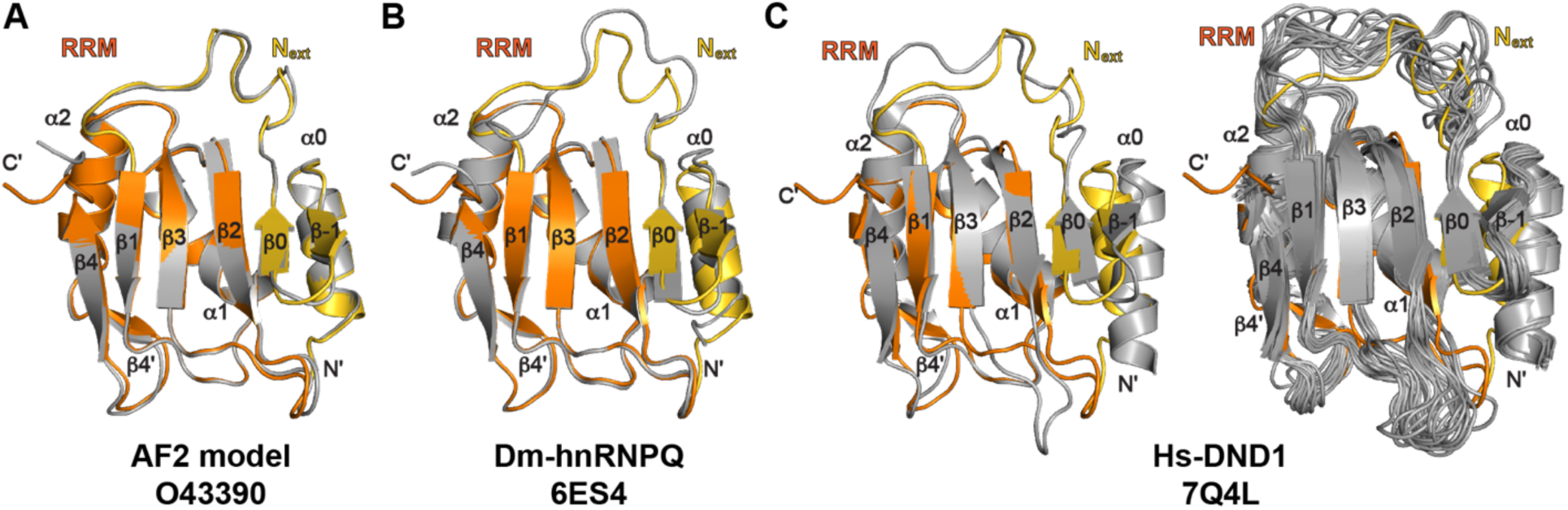
Comparison of predicted and high-resolution structures of eRRM1 domains in the hnRNPR-like family. A) AF2 model (colored in gray) superimposed on X-ray crystal structure of Hs-hnRNPR eRRM1; B) superimposition of Hs-hnRNPR eRRM1 and Dm-hnRNPQ eRRM1 (colored in gray); C) superimposition of Hs-hnRNPR eRRM1 and DND1 (colored in gray), with the lowest energy model shown on the left and the structural ensemble shown on the right.

A solution NMR structural ensemble was reported for the human hnRNPR-like protein DND1 containing eRRM1 and RRM2 domains bound to an RNA oligomer (35). The heavy-atom RMSD between the Hs-hnRNPR eRRM1 and the lowest-energy model in the human DND1 (Hs-DND1) eRRM1 structural ensemble is 1.5 Å for all residues (aa 16-134) and 0.99 Å for structured residues in the RRM (**Figure 6C**). Differences are primarily observed in the N_ext_ motif, particularly helix α0, loop0, and the RRM loopβ2-β3. These differences are likely due to the different method of structure determination and/or the presence of RNA substrate. Inspection of the NMR structural ensemble shows higher RMSD among models in loop0 and loopβ2-β3 regions, suggesting dynamics in these loops consistent with our results for human hnRNPR eRRM1. From comparison of these examples of eRRMs for representative proteins in the hnRNPR-like family, we identify the required elements for the eRRM fold to be the (1) N_ext_ motif, which has an αββ topology; (2) electrostatic interactions between the N_ext_ and RRM; and (3) tryptophan cage motif in loop0.

## Discussion

hnRNPR and hnRNPR-like proteins perform essential roles in RNA processing. However, biophysical and structural information on hnRNPR is sparse, limiting a complete understanding of how hnRNPR achieves its function in RNA transcription and splicing regulation. Here, we determined the X-ray crystal structure of the human hnRNPR N-terminal RRM1 and identified an atypical eRRM consisting of a N_ext_ motif that docks alongside a canonical RRM. We evaluated eRRM1 conformational dynamics using NMR spectroscopy, which showed an overall ordered protein with flexible N- and C-termini. However, these dynamics may differ in the full-length hnRNPR, where the N-terminal end is linked to the NαB domain and the C-terminal end is linked to RRM2. We show that truncation of the N_ext_ motif results in loss of protein solubility. Point substitutions that disrupt N_ext_-RRM interactions result in increased dynamics at the N-terminus and reduced thermal stability. Our work demonstrates that stable association of the N_ext_ motif with the RRM is required for protein solubility and thermal stability. These findings are consistent with previous research that evaluated the impact of N-terminal truncations in the hnRNPR-like protein A1CF. Truncating the first 13 residues at the A1CF N-terminus reduced activity to 33% of full-length A1CF, with complete loss of activity when truncating two additional residues (42). Our data explains this loss of activity due to truncation of the A1CF eRRM1 domain, which we predict to begin at residue 14 (**Figure S1B**).

The eRRM1 is further stabilized by a tryptophan cage motif in loop0, which positions the N_ext_ motif for docking to the β2-α1 side of the RRM. The central tryptophan residue has a dual role in eRRM1 architecture, also participating in the N_ext_-RRM interface. Loop0 residues are highly conserved among hnRNPR proteins and hnRNPR-like family proteins. Using combined NMR spectroscopy and thermal denaturation studies, we show that alanine substitutions to loop0 residues lead to increased local dynamics and reduced thermal stability. Consistent with our results, loop0 residues are highly conserved in hnRNPR proteins, as well as among the hnRNPR-like family. A survey of somatic mutations in cancer studies shows cancer-associated mutations in this loop, suggesting that stable formation of the tryptophan cage in loop0 is necessary for cellular function. The tryptophan cage motif was first identified in the synthetic Trp-cage peptide, and to our knowledge the eRRM represents the first reported example of a tryptophan cage motif in a naturally occurring protein.

RRMs often have additional secondary structure elements that aid in substrate recognition and specificity. Here, the eRRM is a unique example of an extension that is nearly half the size of the RRM and extends the β-sheet surface from five strands to seven strands. The structural basis for hnRNPR eRRM1-RNA recognition remains unknown, and it remains to be seen how the hnRNPR eRRM1 coordinates with the NαB and tandem RRMs to bind RNA and protein substrates. However, a previously determined NMR structure of the human DND1 eRRM1-RRM2 tandem domain bound to RNA provides insights into eRRM1-RNA recognition (35). RNA substrate is sandwiched between eRRM1 and RRM2 domains and binds to the eRRM1 β-sheet surface and loopβ2-β3 (**Figure S18**). Beyond the canonical interactions with RNP1 and RNP2 sequences, RNA substrate also interacts with β2 residue R88 (R195 in hnRNPR) and N_ext_ β-turn residues N37 and Q39 (T141 and Q144 in hnRNPR). We anticipate that the hnRNPR eRRM1 may have a similar mode of RNA recognition in which the β2-strand, β2-β3 loop, and N_ext_ β-hairpin contribute to binding. The DND1 eRRM1 does not interact with loop0 or Y85, the equivalent residue to W192 in hnRNPR. The high sequence conservation of the tryptophan cage suggests a structural role rather than a function in RNA recognition. In summary, this work provides fundamental biophysical and structural insights into the eRRM1 fold to begin to characterize its role in RNA processing and cellular function.

## Methods

### Protein construct design and cloning

The amino acid sequence for human hnRNPR protein (Uniprot ID O43390) was used to generate the gene sequence optimized for *E. coli* codon bias, and purchased as a gBlock (Integrated DNA Technologies, IDT). The gene was cloned into a pET vector containing an N-terminal His-tag, a TEV protease recognition site, and a kanamycin resistance gene. Forward and reverse primers, designed using Benchling and ordered from IDT, were used to clone protein constructs. The plasmids were then transformed into DH5α cells for propagation. Afterward, the plasmids were purified from the DH5α cells using a miniprep plasmid cleanup kit (Zymo Research). Whole Plasmid Sequencing was performed by Plasmidsaurus™ using Oxford Nanopore Technology with custom analysis and annotation. A complete list of protein sequences used in this study is provided in **Table S2**.

### Protein expression and purification

Plasmids were transformed into *E. coli* NiCo21(DE3) competent cells (New England Biolabs). Cells were cultured with shaking in LB broth media with 0.05 mg/mL kanamycin at 37 °C to an OD600 of 0.6. Culture was transferred to 18 °C, expression induced with 0.5 mM isopropyl β-D-1-thiogalactopyranoside (IPTG), and grown for 18–20 h with shaking. For NMR experiments, cells were cultured in M9 media and supplemented with ^13^C-labelled glucose (Cambridge Isotope Laboratories) and/or ^15^N-labelled ammonium chloride (Cambridge Isotope Laboratories). Cells were harvested by centrifugation at 6000 xg for 15 minutes. Cells were resuspended in resuspension buffer (20 mM Na_2_HPO_4_/NaH_2_PO_4_, 1.5 M NaCl, 20 mM imidazole, 10% glycerol, 1 mM tris(2-carboxyethyl) phosphine (TCEP), pH 7.0). 0.1 mM PMSF and 1 mg lysozyme were added prior to sonication to lyse cells. After sonication, soluble and insoluble fractions were separated by centrifugation at 36,000 xg for 45 min, and the supernatant was filtered using a 0.45 μm syringe filter. Affinity purification was performed using nickel nitriloacetic acid (Ni-NTA) affinity column (Qiagen) attached to an AKTA™ start system (Cytiva). After loading the supernatant, the column was washed with 10 column volumes (CV) of resuspension buffer followed by elution in a linear gradient with elution buffer (20 mM Na_2_HPO_4_/NaH_2_PO_4_, 300 mM NaCl, 1.0 M imidazole, 10% glycerol, 1 mM TCEP, pH 7.0). Fractions were analyzed by SDS-PAGE 4–12% Bis-Tris NuPAGE™ gels (ThermoScientific). To remove the His-tag, protein was dialyzed against storage buffer (20 mM Na_2_HPO_4_/NaH_2_PO_4_, 150 mM NaCl, 1 mM TCEP, pH 6.0) in the presence of TEV protease (Addgene #92414, recombinantly expressed and purified in-house) (43) for 3-4 hrs at room temperature or overnight at 4 °C. After dialysis, cleaved protein was filtered using a 0.45 μm syringe filter and purified by a second Ni–NTA purification step (Qiagen). As a final purification step, proteins were purified by size exclusion chromatography using a HiLoad16/600 Superdex 75 pg (Cytiva) column attached to an AKTA™ Pure M25 FPLC system (Cytiva) in storage buffer. Fractions were analyzed by 4–12% Bis-Tris NuPAGE™ gel and inspection A260/A280 ratios to identify fractions with pure protein. Protein concentration was determined from the absorbance measured at 280 nm using a NanoDrop One™ spectrophotometer (Thermo Fisher Scientific) using Beer’s Law. Pure fractions were concentrated to 0.1-0.8 mM using a 3 kDa Amicon™. Expasy ProtParam (44) was used to compute protein extinction coefficients, molecular weights, and theoretical pI values.

### X-ray crystallography

Proteins were crystalized using the hanging drop vapor diffusion method. Proteins were prepared in crystallization buffer (20 mM HEPES, 150 mM NaCl, 1 mM TCEP pH = 7.0) at 16 mg/mL. Rod-like crystals appeared within 48 hours and were harvested at 200-300 μm in size. Crystals formed under reservoir conditions of 0.04 M monobasic potassium phosphate, 20% (v/v) glycerol, and 8% (w/v) polyethylene glycol 8000 with a 1:1 protein-to-reservoir ratio. Data was collected at 100 K at the Stanford Synchrotron Radiation Lightsource at the SLAC National Accelerator Laboratory on beamline 12-2 using 0.72929 Å X-rays with a Dectris Pilatus 6M detector. Data from a single crystal was used for structure determination. Data was indexed and integrated using XDS (45). Data reduction was performed using Aimless, Pointless, and Ctruncate in the CCP4 suite (46). Phases were determined using molecular replacement against the X-ray crystal structure of the drosophila hnRNPQ eRRM (PDB ID 6ES4) (34). Model building and refinement was performed using Coot version 0.9.8.94 (47) and PHENIX version 1.21.1 (48) with TLS refinement (49). Final data collection, phasing, and refinement statistics are reported in **Table 1**.

### NMR spectroscopy

NMR experiments were performed at 25 °C on a 600 MHz Bruker spectrometer equipped with a triple-resonance HCN cryoprobe and Avance Neo console. NMR samples were prepared in storage buffer with added 5% D_2_O at 0.3-0.8 mM concentrations in 3 mm or 5 mm NMR tubes (Norell). Backbone (N, H, Cα, Cβ, CO) resonance assignments were performed using standard triple resonance experiments (50; 51). Data were processed using NMRPipe (36) and analyzed using NMRFAM-Sparky 1.470 (52) in the NMRbox virtual machine (53).

^1^H-^15^N heteronuclear nuclear Overhauser effect (NOE) experiments (hsqcnoef3gpsi) from the Bruker experimental suite were recorded in an interleaved manner with 32 scans and 2 s incremental delay for 0.5 mM protein samples. The heteronuclear NOE is reported as the residue-specific ratio of peak intensity between the saturated and unsaturated experiments (I_sat_/I_unsat_). Error was estimated as the standard deviation of noise in the saturated (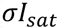) and unsaturated (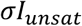) experiments (54; 55): 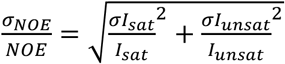. R_1_ and R_2_ values were obtained from T_1_ (hsqct1etf3gpsi3d) and T_2_ (hsqct2etf3gpsi3d) relaxation experiments with 32 scans and 2 s incremental delay. Data was processed in NMRPipe (36) and relaxation rates were calculated using the Function and Data Analysis (FuDA) software package (56). For T_1_ experiments, delays were 20ms, 60ms x 2, 200 ms, 400 ms, 800 ms x 2, and 1200 ms. T_2_ loops (2 x 2, 4, 6, 8, 12 x 2, 16) had a 0.01696 ms variable delay. Weighted average chemical shift perturbations were calculated using the equation 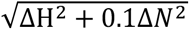 (51).

### Circular dichroism (CD)

Thermal denaturation experiments were performed on a Jasco815 spectrometer equipped with a Peltier temperature control device. Protein samples were prepared in storage buffer at concentrations ranging between 18–25 μM. CD spectra were collected at 222 nm with the following parameters: temperature range of 10–90 °C (except for the W192A construct, where the temperature range used was 4–80 °C), 1 sec hold time, 1.0 °C/sec ramp rate, 1 °C sampling interval, 5 sec wait time, 1 sec data integration time, 2 nm band width, and standard sensitivity. Data were analyzed using in-house python code (https://github.com/eichhorn-lab). Due to protein aggregation at temperatures above 70 °C, data from 10 °C-70 °C (except for W192A, which included data from 4 °C-70°C), were normalized using the equation (*data point* − *min. value*)(*max. value − min. value*) to compute percent folded protein. The data was fitted with a logistic function of 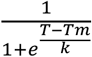, where Tm is the melting temperature and k is the slope factor. Independent experiments were performed in duplicate. A complete list of fitted parameters and replicates for each construct is provided in **Table S3**.

### Bioinformatics

The human hnRNPR sequence (O43390) was retrieved from the UniProt database (57) and used as a query in UniProt BLAST (58) with default search parameters. Of 250 initial hits, 25 were excluded to remove redundant entries, achieved sequences, and multiple isoforms from the same species. The selected homologous sequences were then aligned using the EMBL Clustal Omega Multiple Sequence Alignment (MSA) tool (59; 60). The alignment quality was visually inspected to ensure accurate residue mapping. To assess the evolutionary conservation of individual residues, the MSA of the full-length hnRNPR was analyzed using the ConSurf web server (61; 62) using default parameters. Conservation scores were mapped onto the AF2 predicted model (AF-O43390-F1). For cancer-related missense mutation analysis, data was downloaded from the COSMIC database (40; 41) (Supplementary File 1). An in-house python script was used to extract missense mutations; sort data by cancer histology subtype, primary tissue, and domain location; and generate plots (https://github.com/eichhorn-lab).

## Supporting information

hnRNPR Missense mutations from COSMIC database

hnRNPR MSA gapped fasta file

## Supplementary Material

Supplementary tables and figures, the fasta file of the MSA of hnRNPR proteins, and the excel spreadsheet containing missense mutations identified from COSMIC data have been included as supplementary material.

## Data availability

PDB coordinates are deposited to the PDB with ID XXXX. NMR resonance assignments, R_1_ and R_2_ relaxation, and ^1^H-^15^N NOE data are deposited to BMRB ID xxxxxx. Python scripts are deposited to github (https://github.com/eichhorn-lab).

## Declaration of competing interest

The authors declare no competing interest.

## Acknowledgements

We acknowledge funding support from the National Institutes of Health (1R35GM143030) and the Nebraska Center for Integrated Biomolecular Communication (P20 GM113126). Use of the Stanford Synchrotron Radiation Lightsource, SLAC National Accelerator Laboratory, is supported by the U.S. Department of Energy, Office of Science, Office of Basic Energy Sciences under Contract No. DE-AC02-76SF00515. The SSRL Structural Molecular Biology Program is supported by the DOE Office of Biological and Environmental Research, and by the National Institutes of Health, National Institute of General Medical Sciences (P30GM133894). The contents of this publication are solely the responsibility of the authors and do not necessarily represent the official views of NIGMS or NIH. We acknowledge assistance from Linh Hua in buffer preparation. We thank all members of the Eichhorn lab for critical reading of the manuscript and providing constructive feedback.

## Notes

### Competing Interest Statement

The authors have declared no competing interest.

