## Supplementary File 1 for "An evolutionarily conserved tryptophan cage promotes folding of the extended RNA recognition motif in the hnRNPR-like protein family"

**List of supplemental tables and figures:**

**Table S1:** Crystallography statistics for *H. sapiens* hnRNPR eRRM1
**Table S2:** Protein constructs used in this study
**Table S3:** Fitted parameters from circular dichroism (CD) melting experiments
**Figure S1**: hnRNPR-like protein domain topology and predicted structure
**Figure S2:** NMR and CD analysis of WT eRRM1 structure and thermal stability
**Figure S3:** Secondary structure prediction from TALOS+ chemical shift analysis
**Figure S4:** Sequence conservation of eRRM1 in verified hnRNPR proteins
**Figure S5:** Sequence conservation of verified and predicted hnRNPR proteins
**Figure S6:** SDS-PAGE gel of RRM1 (aa162-246) samples during tandem purification
**Figure S7:** NMR and CD analysis of Ext structure and thermal stability
**Figure S8:** NMR and CD analysis of R133A structure and thermal stability
**Figure S9:** $^{1}$H-$^{15}$N HSQC spectrum of arginine N$\varepsilon$H$\varepsilon$ resonances
**Figure S10:** NMR and CD analysis of Y177A structure and thermal stability
**Figure S11:** NMR and CD analysis of R195A structure and thermal stability
**Figure S12:** NMR and CD analysis of Y156A structure and thermal stability
**Figure S13:** NMR and CD analysis of P161A structure and thermal stability
**Figure S14:** NMR and CD analysis of P150A structure and thermal stability
**Figure S15:** NMR and CD analysis of P151A structure and thermal stability
**Figure S16:** NMR and CD analysis of W192A structure and thermal stability
**Figure S17:** Cancer-associated somatic mutations in hnRNPR
**Figure S18:** RNA recognition of DND1 eRRM1

**Table S1. Crystallography statistics for *H. sapiens* hnRNPR eRRM1**

| | |
|---|---|
| **Data Collection** | |
| Wavelength (Å) | 0.72929 |
| Space Group | P31 2 1 |
| Cell dimensions | |
| a, b, c (Å) | 77.023, 77.023, 52.893 |
| α, β, γ (°) | 90.0, 90.0, 120.0 |
| Resolution (Å) | 1.90 |
| $R_{merge}$ | 0.033 (2.174) |
| $R_{measure}$ | 0.035 (2.281) |
| $R_{pim}$ | 0.011 (0.686) |
| I/σ | 40.7 (1.5) |
| Completeness (%) | 100 (99.9) |
| No. of total reflections | 304500 |
| No. of unique reflections | 14604 |
| Multiplicity | 20.9 (21) |
| Wilson B factor (Å$^2$) | 49.44 |
| $CC_{1/2}$ | 1.000 (0.626) |
| **Refinement** | |
| Resolution (Å) | 31.13-1.90 |
| No. of reflections | 14582 |
| $R_{work}/R_{free}$ | 0.1966/0.2266 |
| No. of atoms | 1042 |
| Protein | 972 |
| Water | 70 |
| Average B factors (Å$^2$) | 63.16 |
| Protein | 62.96 |
| Water | 65.96 |
| Clashscore | 1.03 |
| r.m.s.d | |
| Bond lengths (Å) | 0.007 |
| Bond angles (°) | 0.92 |
| Ramachandran plot | |
| Favored (%) | 99.20 |
| Allowed (%) | 0.80 |
| Outliers (%) | 0.00 |

**Table S2. Protein constructs used in this study**

| Construct | Amino acid boundary (aa) | Amino acid residues (N' – C') | Extinction coefficient ($M^{-1}cm^{-1}$) |
|---|---|---|---|
| WT | 121-246 | MGSSHHHHHHQENLYFQSGPDEAKIKALLERTGY<br>TLDVTTGQRKYGGPPPDSVYSGVQPGIGTEVFV<br>GKIPRDLYEDELVPLFEKAGPIWDLRLMMDPLSG<br>QNRGYAFITFCGKEAAQEAVKLCDSYEIRPGKHL<br>GVCISVANN | 15930 |
| **N-terminal extension and truncation constructs** | | | |
| Ext | 116-246 | MGSSHHHHHHQENLYFQS**QESTK**GPDEAKIKALL<br>ERTGYTLDVTTGQRKYGGPPPDSVYSGVQPGIG<br>TEVFVGKIPRDLYEDELVPLFEKAGPIWDLRLMM<br>DPLSGQNRGYAFITFCGKEAAQEAVKLCDSYEIR<br>PGKHLGVCISVANN | 15930 |
| RRM1 | 162-246 | MGSSHHHHHHQENLYFQSQSGIGTEVFVGKIPRD<br>LYEDELVPLFEKAGPIWDLRLMMDPLSGQNRGY<br>AFITFCGKEAAQEAVKLCDSYEIRPGKHLGVCISV<br>ANN | 11460 |
| **$N_{ext}$-RRM1 interface point substitution constructs** | | | |
| R133A | 121-246 | MGSSHHHHHHQENLYFQSGPDEAKIKALLE**A**TGY<br>TLDVTTGQRKYGGPPPDSVYSGVQPGIGTEVFV<br>GKIPRDLYEDELVPLFEKAGPIWDLRLMMDPLSG<br>QNRGYAFITFCGKEAAQEAVKLCDSYEIRPGKHL<br>GVCISVANN | 15930 |
| R145A | 121-246 | MGSSHHHHHHQENLYFQSGPDEAKIKALLERTGY<br>TLDVTTGQ**A**KYGGPPPDSVYSGVQPGIGTEVFV<br>GKIPRDLYEDELVPLFEKAGPIWDLRLMMDPLSG<br>QNRGYAFITFCGKEAAQEAVKLCDSYEIRPGKHL<br>GVCISVANN | 15930 |
| R195A | 121-246 | MGSSHHHHHHQENLYFQSGPDEAKIKALLERTGY<br>TLDVTTGQRKYGGPPPDSVYSGVQPGIGTEVFV<br>GKIPRDLYEDELVPLFEKAGPIWDL**A**LMMDPLSG<br>QNRGYAFITFCGKEAAQEAVKLCDSYEIRPGKHL<br>GVCISVANN | 15930 |
| Y177A | 121-246 | MGSSHHHHHHQENLYFQSGPDEAKIKALLERTGY<br>TLDVTTGQRKYGGPPPDSVYSGVQPGIGTEVFV<br>GKIPRDL**A**EDELVPLFEKAGPIWDLRLMMDPLSG<br>QNRGYAFITFCGKEAAQEAVKLCDSYEIRPGKHL<br>GVCISVANN | 14440 |
| **Loop0 point substitution constructs** | | | |
| P150A | 121-246 | MGSSHHHHHHQENLYFQSGPDEAKIKALLERTGY<br>TLDVTTGQRKYGG**A**PPDSVYSGVQPGIGTEVFV<br>GKIPRDLYEDELVPLFEKAGPIWDLRLMMDPLSG<br>QNRGYAFITFCGKEAAQEAVKLCDSYEIRPGKHL<br>GVCISVANN | 15930 |
| P151A | 121-246 | MGSSHHHHHHQENLYFQSGPDEAKIKALLERTGY<br>TLDVTTGQRKYGGP**A**PDSVYSGVQPGIGTEVFV | 15930 |

| | | | |
|---|---|---|---|
| | | GKIPRDLYEDELVPLFEKAGPIWDLRLMMDPLSG<br>QNRGYAFITFCGKEAAQEAVKLCDSYEIRPGKHL<br>GVCISVANN | |
| Y156A | 121-246 | MGSSHHHHHQENLYFQSGPDEAKIKALLERTGY<br>TLDVTTGQRKYGGPPPDSV**A**SGVQPGIGTEVFV<br>GKIPRDLYEDELVPLFEKAGPIWDLRLMMDPLSG<br>QNRGYAFITFCGKEAAQEAVKLCDSYEIRPGKHL<br>GVCISVANN | 14440 |
| P161A | 121-246 | MGSSHHHHHQENLYFQSGPDEAKIKALLERTGY<br>TLDVTTGQRKYGGPPPDSVYSGVQ**A**GIGTEVFV<br>GKIPRDLYEDELVPLFEKAGPIWDLRLMMDPLSG<br>QNRGYAFITFCGKEAAQEAVKLCDSYEIRPGKHL<br>GVCISVANN | 15930 |
| W192A | 121-246 | MGSSHHHHHQENLYFQSGPDEAKIKALLERTGY<br>TLDVTTGQRKYGGPPPDSVYSGVQPGIGTEVFV<br>GKIPRDLYEDELVPLFEKAGPI**A**DLRLMMDPLSG<br>QNRGYAFITFCGKEAAQEAVKLCDSYEIRPGKHL<br>GVCISVANN | 10430 |

Nonnative sequence used for affinity purification and TEV cleavage are indicated in gray. Residues that differ from WT are indicated in bold and underlined. Extinction coefficient was estimated for each construct using the Protparam webserver tool (1).

**Table S3. Fitted parameters from circular dichroism (CD) melting experiments**

| Construct | Protein boundary (aa) | Tm (°C) | Slope factor (k) | Replicates |
|---|---|---|---|---|
| WT | 121-246 | 59 ± 1 | 1 ± 0.1 | 4 |
| **N-terminal extension and truncation constructs** | | | | |
| Ext | 116-246 | 62 ± 1 | 3 ± 0.2 | 4 |
| RRM1 | 162-246 | ND | ND | ND |
| **$N_{ext}$-RRM1 interface point substitution constructs** | | | | |
| R133A | 121-246 | 55 ± 1 | 1 ± 0.1 | 3 |
| R145A | 121-246 | ND | ND | ND |
| R195A | 121-246 | 50 ± 1 | 3 ± 0.3 | 2 |
| Y177A | 121-246 | 47 ± 0.1 | 2 ± 0.1 | 3 |
| **Loop0 point substitution constructs** | | | | |
| P150A | 121-246 | 49 ± 0.9 | 2 ± 0.1 | 3 |
| P151A | 121-246 | 46 ± 0.2 | 1 ± 0.2 | 2 |
| Y156A | 121-246 | 57 ± 1 | 1 ± 0.3 | 2 |
| P161A | 121-246 | 52 ± 0.7 | 1 ± 0.2 | 3 |
| W192A | 121-246 | 40 ± 0.2 | 6 ± 0.4 | 5 |

ND due to insoluble expression

Slope factor (k) describes the cooperativity of the melting transition, with low values indicating more cooperative melting and high values indicating less cooperative melting.

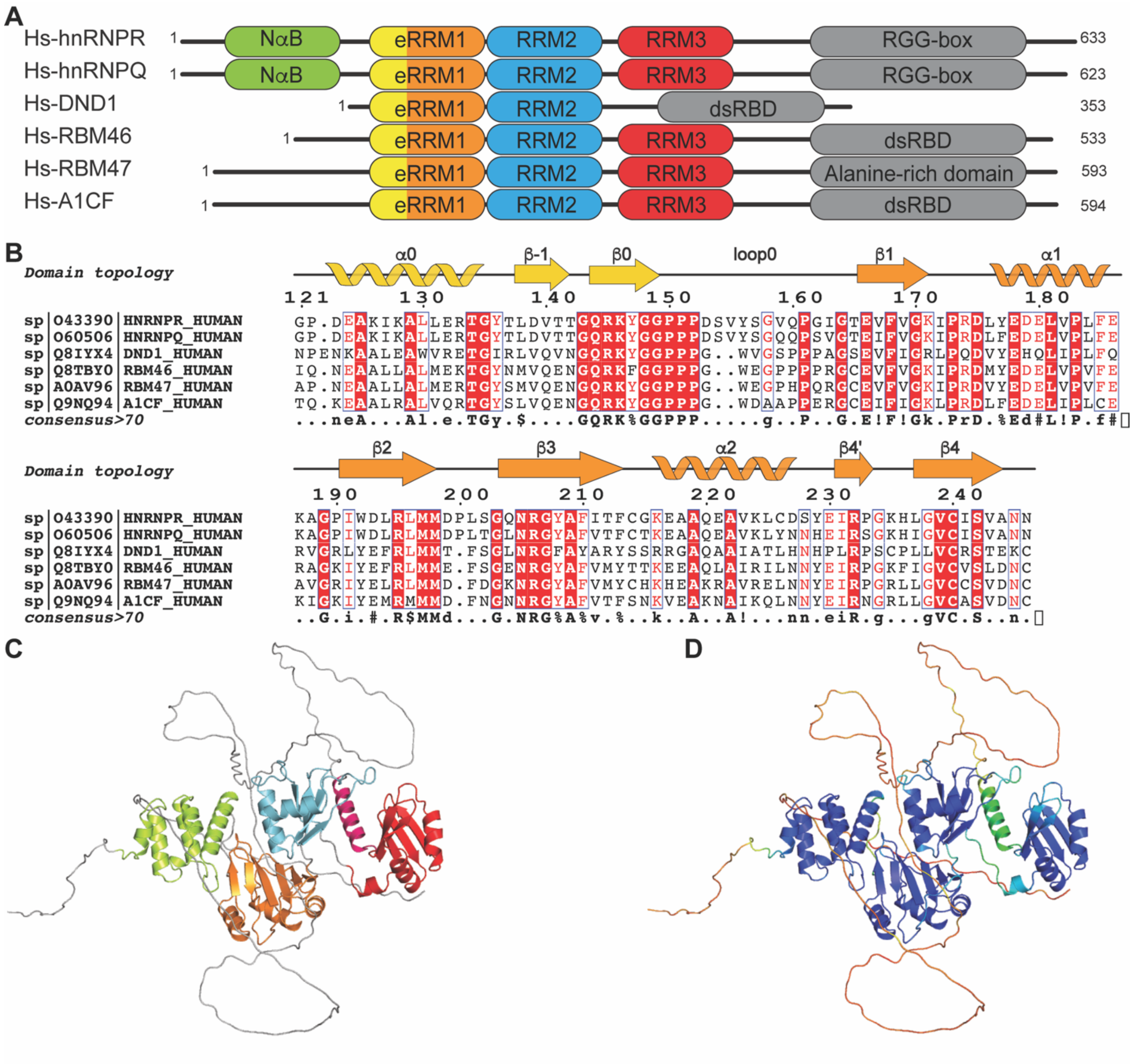

**Figure S1.** hnRNPR-like protein domain topology and predicted structure. A) The domain topology of hnRNPR-like proteins reveals a conserved eRRM1 domain. All proteins contain at least two tandem RRMs, with the eRRM1 located near the N-terminus and as the first RRM in the tandem RRM arrangement; B) Multiple sequence alignment (MSA) of the eRRM1 domain across human hnRNPR-like family proteins reveals high sequence conservation within loop0 and β2, both of which are critical for forming the tryptophan cage motif. C-D) AlphaFold2 model of human hnRNPR (Uniprot ID O43390) C) colored by domain where NαB is colored lime, eRRM1 is colored orange, RRM2 is colored light blue, RRM3 is colored red, and disordered regions are colored gray. D) colored by AF2-computed confidence in structure prediction, where blue is most confident and red is least confident.

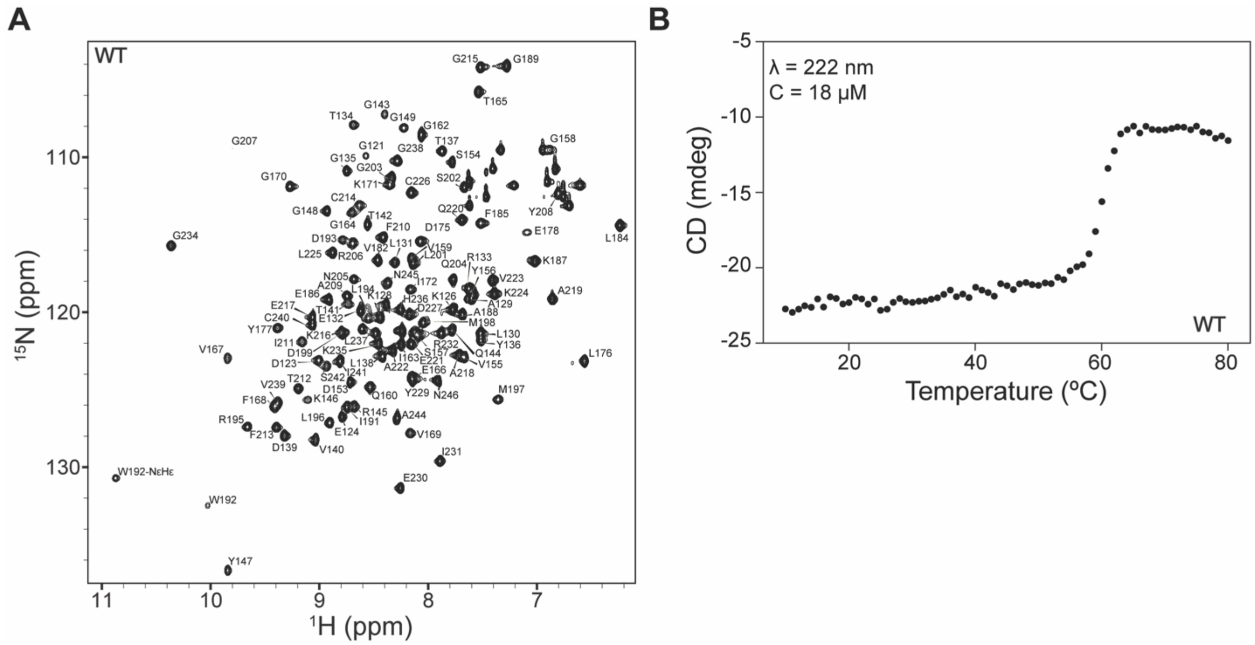


**Figure S2.** NMR and CD analysis of WT eRRM1 structure and thermal stability. A) $^{1}$H-$^{15}$N HSQC spectrum showing backbone amide resonance assignments of WT; B) A representative plot of the raw CD melting trace showing a reduction in molar ellipticity as a function of temperature indicating unfolding of WT. Legend: λ is the wavelength at which data was collected, C is the concentration of the protein sample.

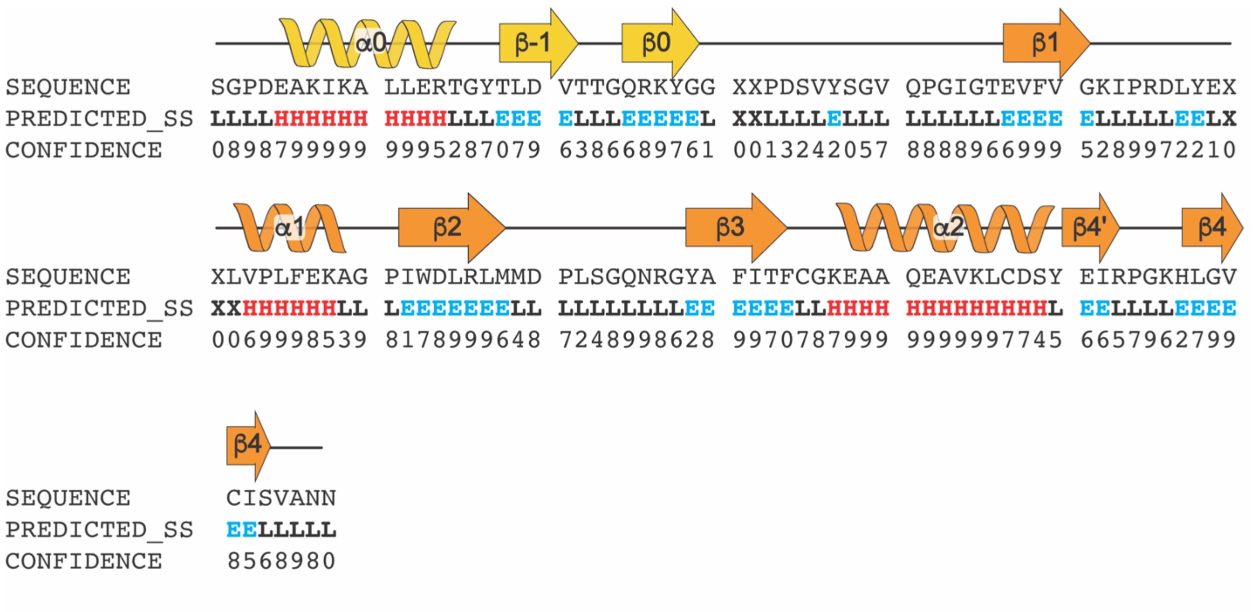


**Figure S3.** Secondary structure prediction from TALOS+ chemical shift analysis. Diagram indicates secondary structure observed in X-ray crystal structure, with $N_{ext}$ colored gold and RRM colored orange. Residues predicted to be random coil are labeled L, α-helical are labeled H and colored red, and β-strand are labeled E and colored blue. TALOS+ confidence in the prediction is indicated on a scale of 0-9, where 9 is most confident. The predicted secondary structure is consistent with the experimentally determined X-ray crystallographic structure.

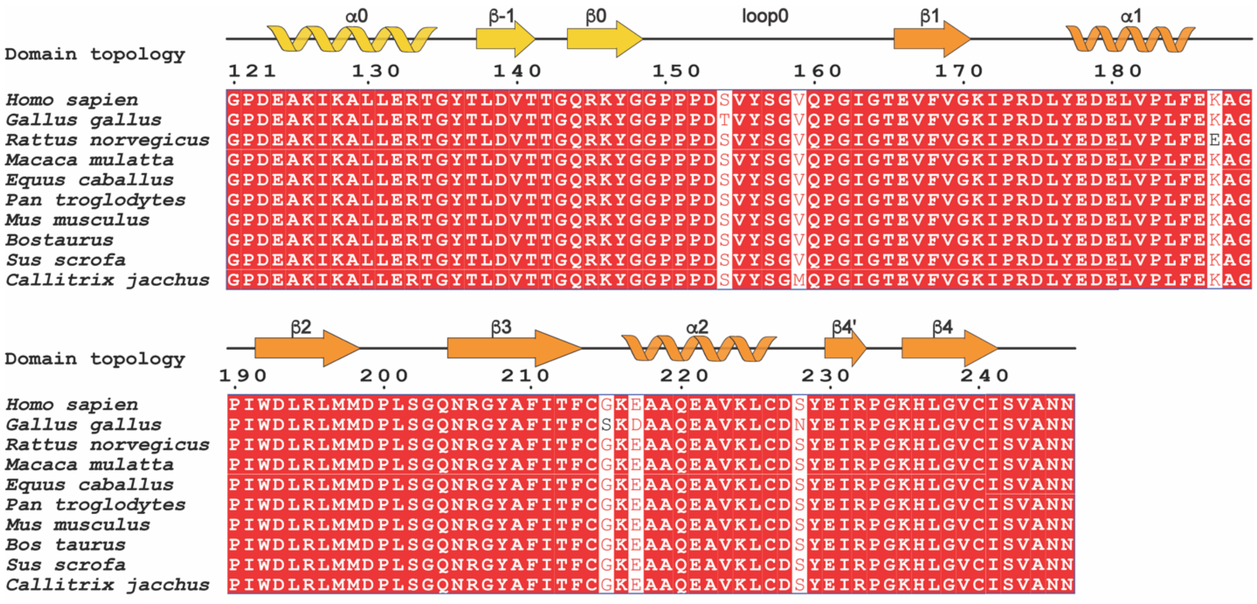


**Figure S4.** Sequence conservation of eRRM1 in verified hnRNPR proteins. MSA of the eRRM1 domains of verified hnRNPR proteins across ten species demonstrates high sequence conservation.

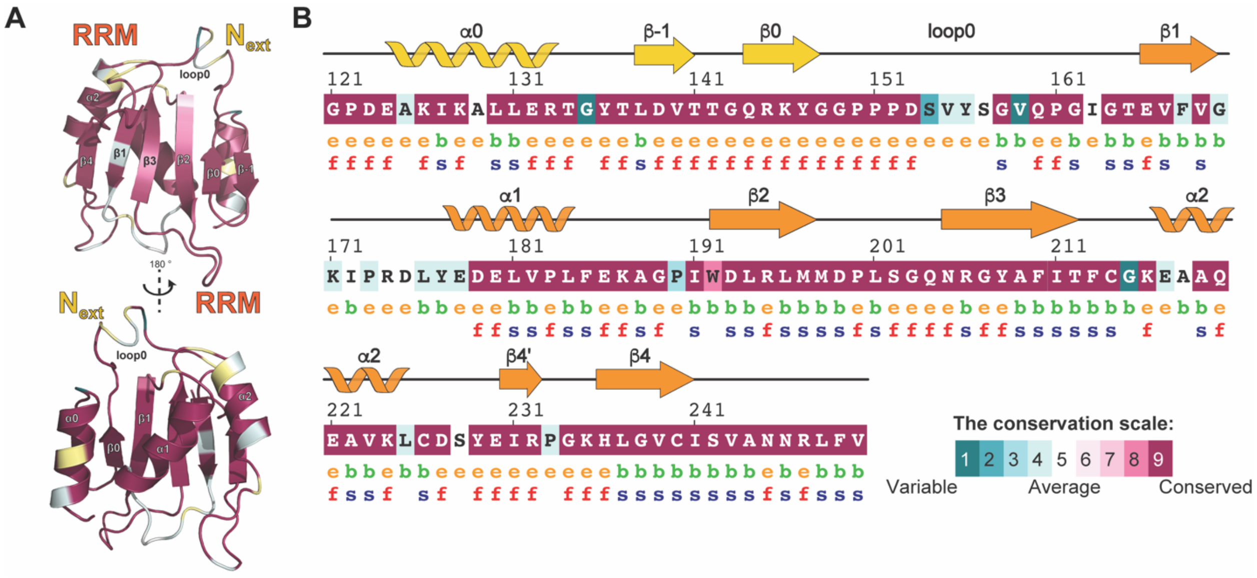


**Figure S5:** Sequence conservation of eRRM1 in verified and predicted hnRNPR proteins. A) Consurf conservation scale mapped onto the AF2 predicted model of human hnRNPR eRRM1 domain for both predicted and verified hnRNPR proteins; B) human hnRNPR eRRM1 sequence colored by Consurf conservation scale on a scale of 1-9, where maroon is most conserved and teal is most variable. Residues predicted to be solvent exposed are labeled e and colored orange, and those predicted to be buried are labeled b and colored green. Residues predicted to be functional are labeled f and colored red, and predicted to be structural are labeled s and colored blue.

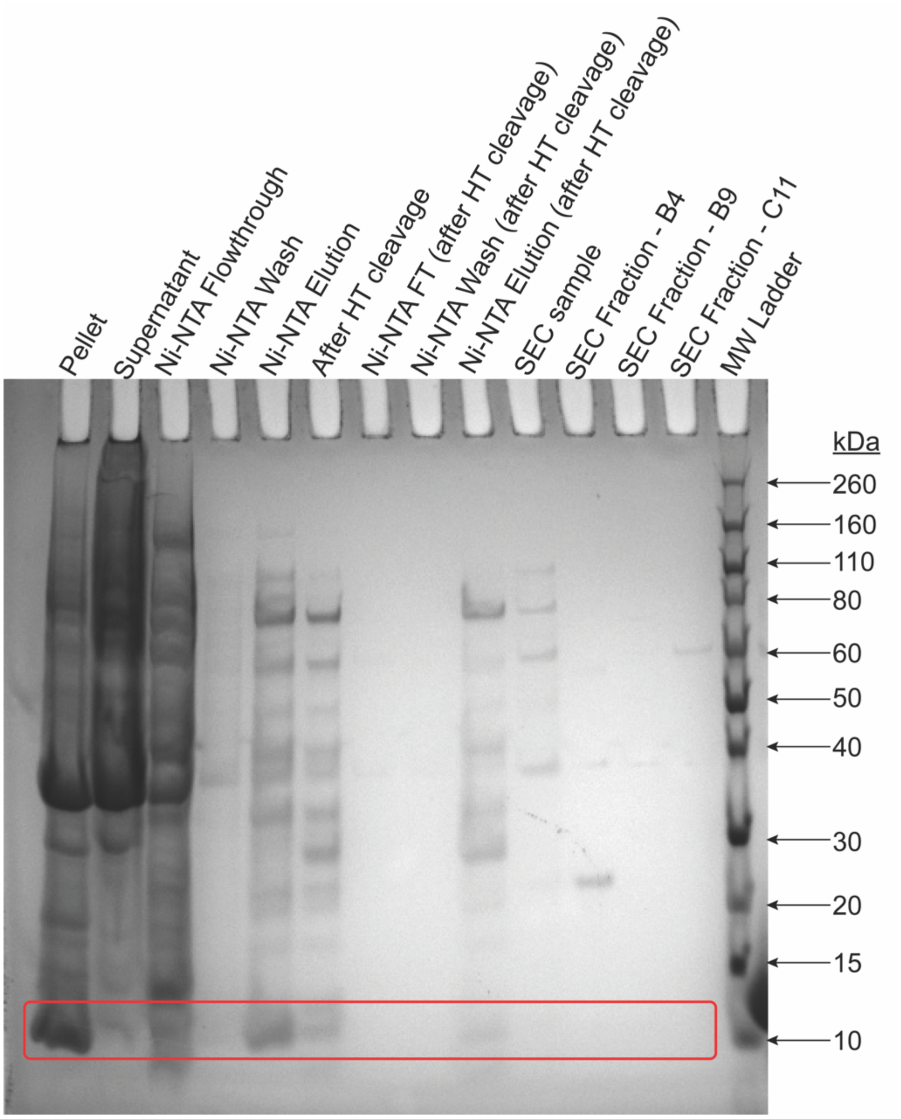


**Figure S6.** SDS-PAGE gel of RRM1 (aa162-246) samples during tandem purification. RRM1 protein construct is largely found in the insoluble fraction, indicated in a red box. Ni_NTA = nickel-charged affinity column purification, HT = His-tag, FT = flowthrough, SEC = size exclusion column chromatography purification, MW = molecular weight.

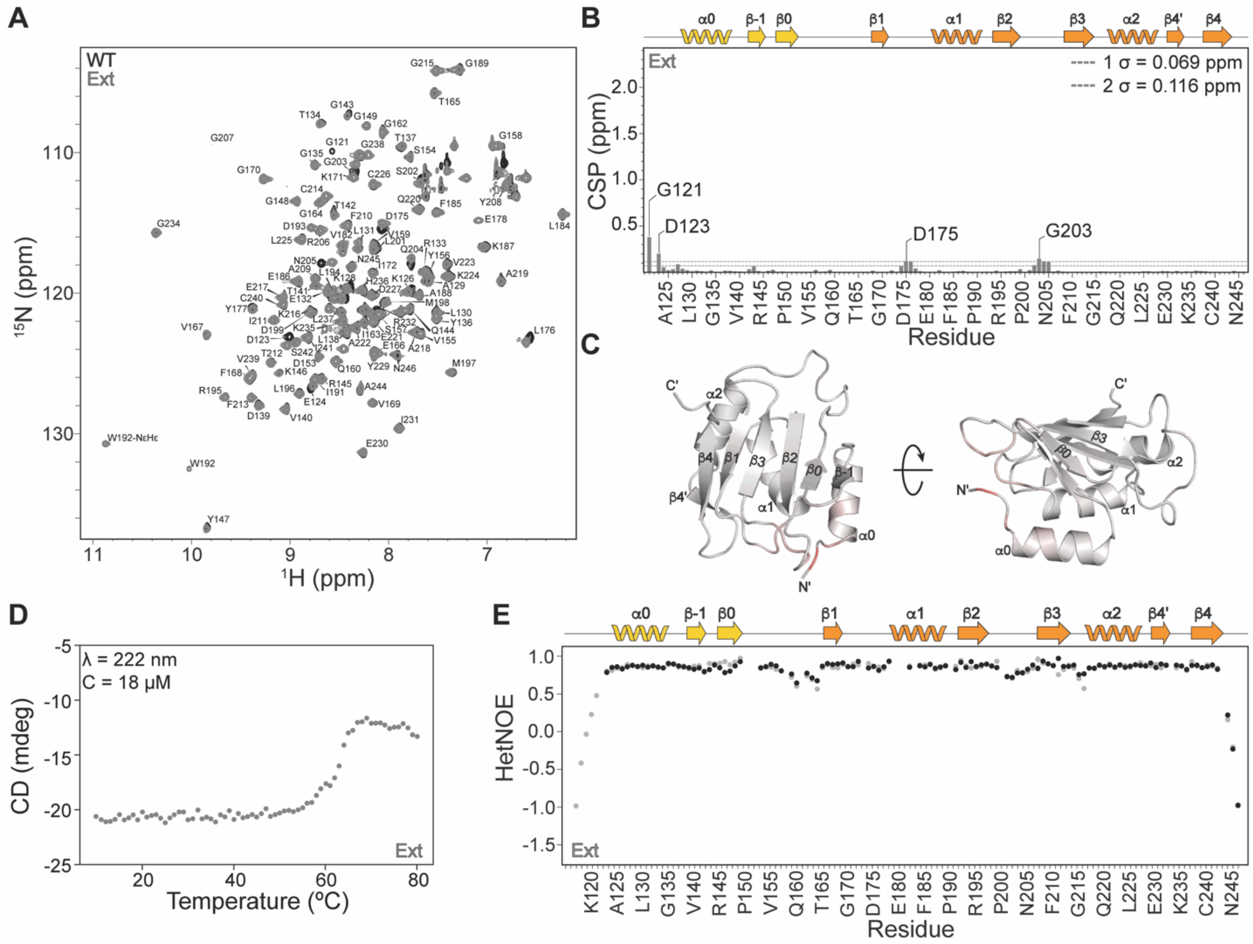


**Figure S7.** NMR and CD analysis of Ext structure and thermal stability. A) Overlay of $^{1}$H-$^{15}$N HSQC spectra comparing backbone amide resonances of Ext and WT eRRM1 constructs. B-C) Weighted-average chemical shift perturbation (CSP) B) shown as a bar plot per residue, where dashed lines indicate 1σ or 2σ standard deviation, also labeled inset, and C) mapped onto the X-ray crystal structure; D) Representative raw CD melting trace of Ext; E) Overlay of $^{1}$H-$^{15}$N heteronuclear NOE plot of Ext (colored in gray) and WT (colored in black).

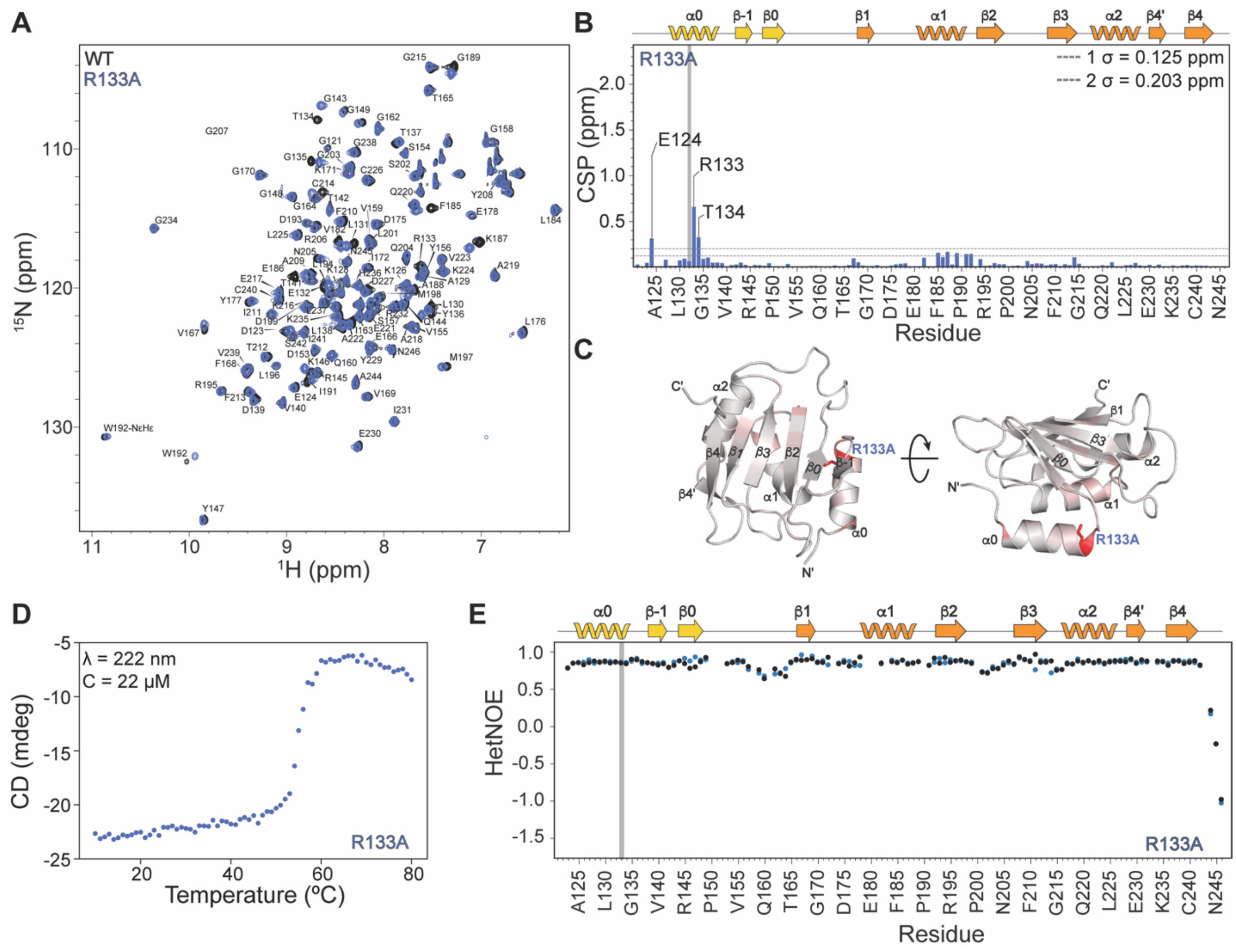


**Figure S8.** NMR and CD analysis of R133A structure and thermal stability. A) Overlay of $^{1}H$-$^{15}N$ HSQC spectra comparing backbone amide resonances of R133A and WT; B) Weighed-average CSP plot shows significant CSPs (>2σ) are localized to substitution site and helix α0; C) CSP values mapped onto the X-ray crystal structure shows most significant CSPs near R133A substitution site, with minor CSPs across the β-sheet; D) Representative raw CD melting trace of R133A; E) Overlay of $^{1}H$-$^{15}N$ heteronuclear NOE plot of R133A (colored blue) and WT (colored black). The substitution site is shown as a gray bar.

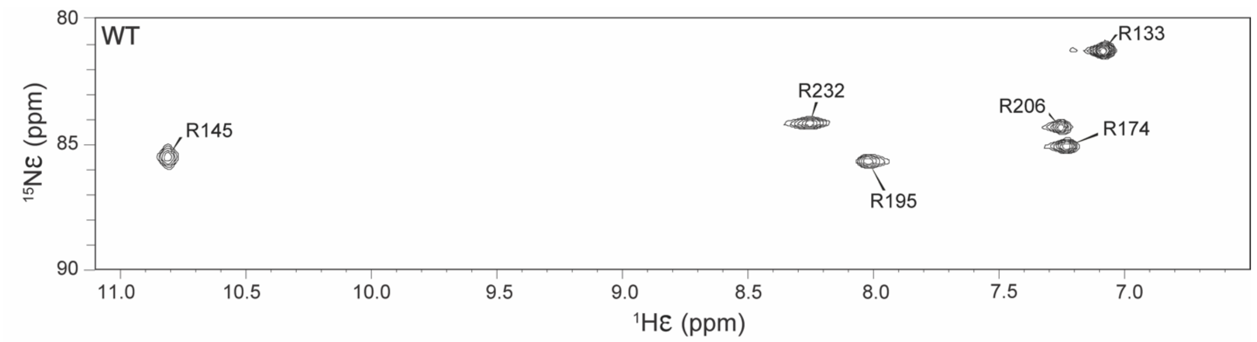


**Figure S9.** $^{1}H$-$^{15}N$ HSQC spectrum of the arginine sidechain (NεHε) resonances of WT. R145 has a distinct downfield shift providing insights into its unique local chemical environment. Resonance assignments were determined from absence of NεHε resonances in alanine substitution constructs.

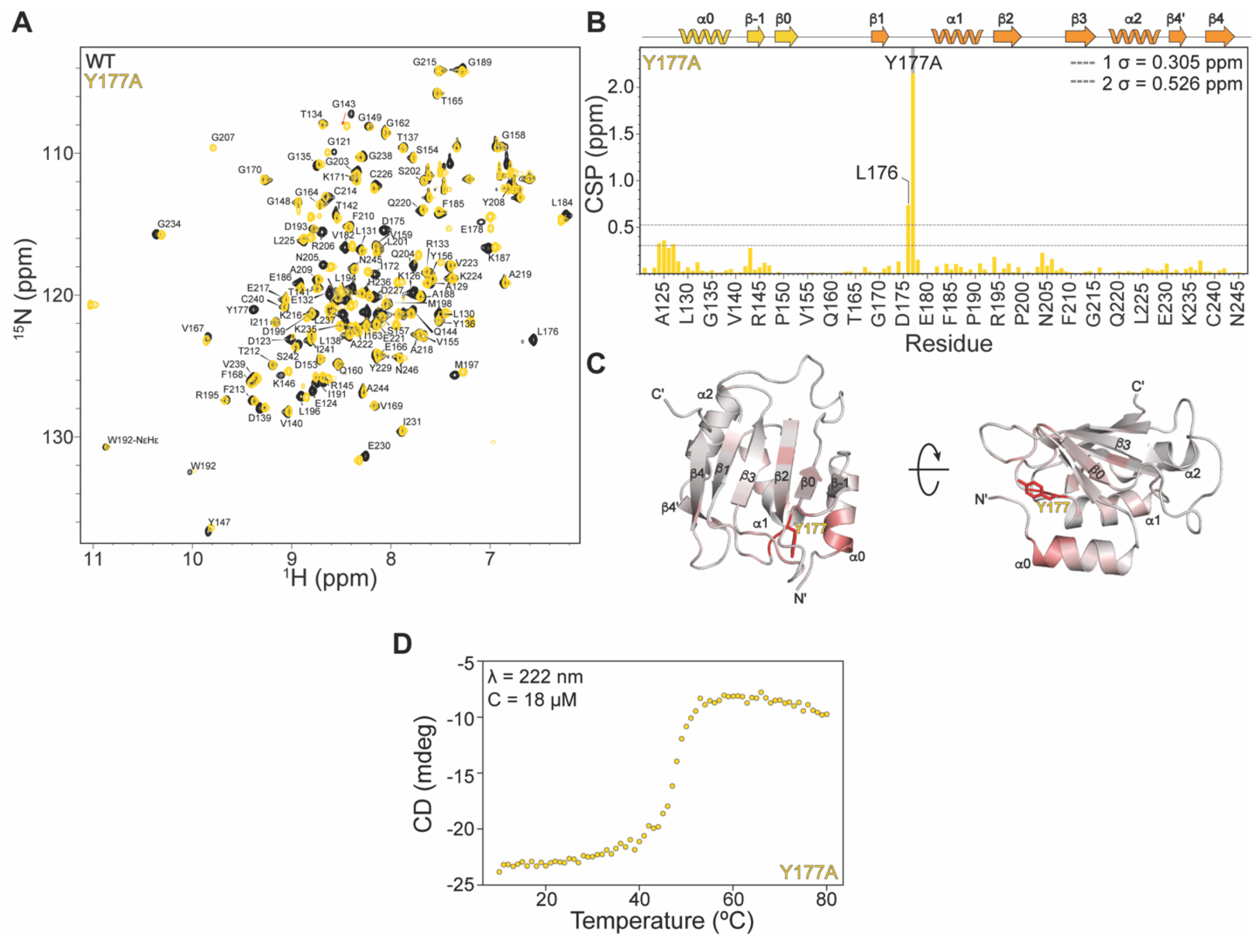


**Figure S10.** NMR and CD analysis of Y177A structure and thermal stability. A) Overlay of $^{1}$H-$^{15}$N HSQC spectra comparing backbone amide resonances of Y177A and WT; B) Plot showing Y177A CSPs; C) CSP values mapped onto the X-ray crystal structure; D) Representative raw CD melting trace of Y177A. The substitution site is shown as a gray bar.

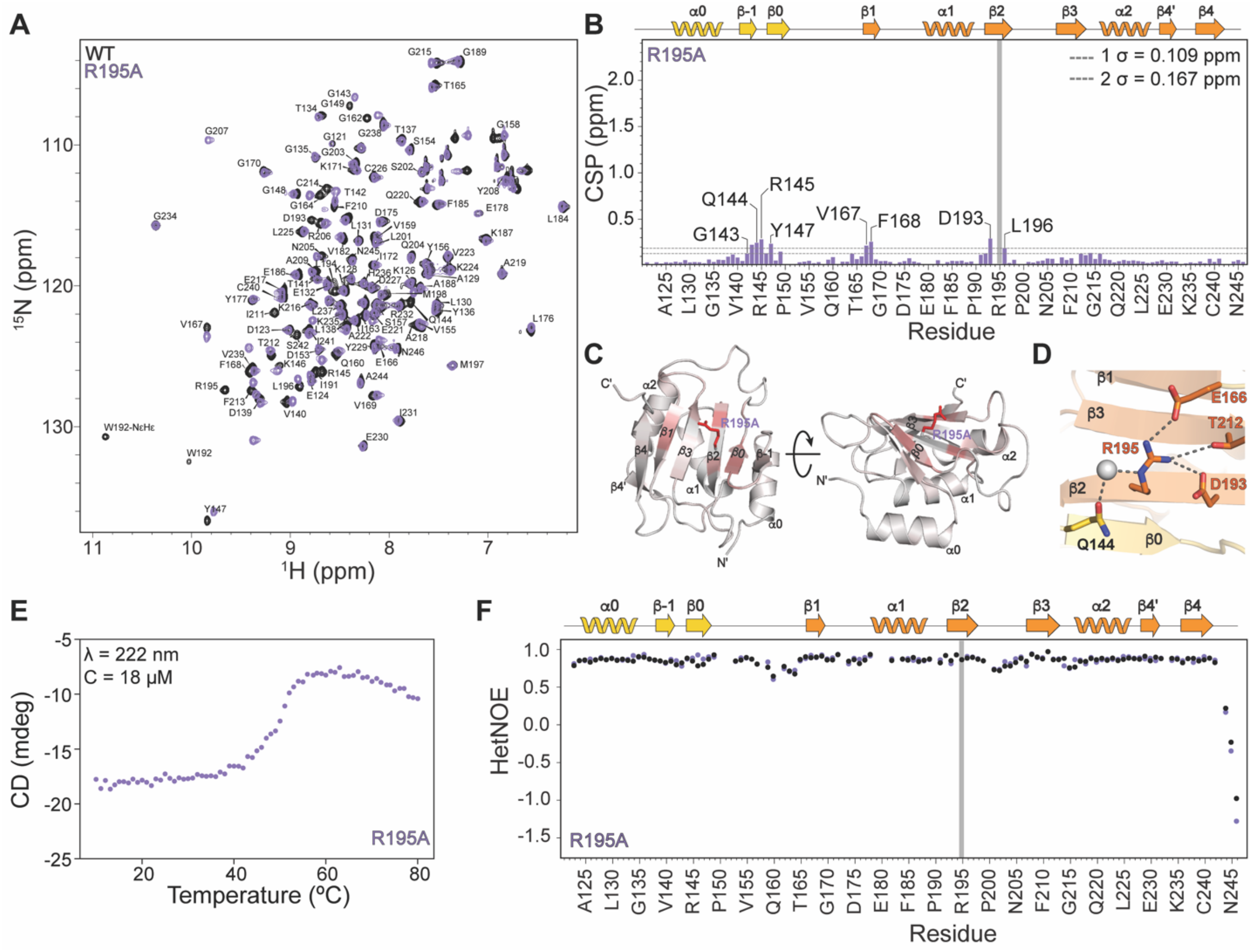


**Figure S11.** NMR and CD analysis of R195A structure and thermal stability. A) Overlay of $^{1}H$-$^{15}N$ HSQC spectra comparing backbone amide resonances of R195A and WT; B) Plot of R195A CSPs; C) R195A CSP values mapped onto the X-ray crystal structure; D) Residue R195 interaction network across the β-sheet surface; E) Representative raw CD melting trace of R195A; F) Overlay of $^{1}H$-$^{15}N$ heteronuclear NOE plot of R133A (colored in purple) and WT (colored in black). The substitution site is shown as a gray bar.

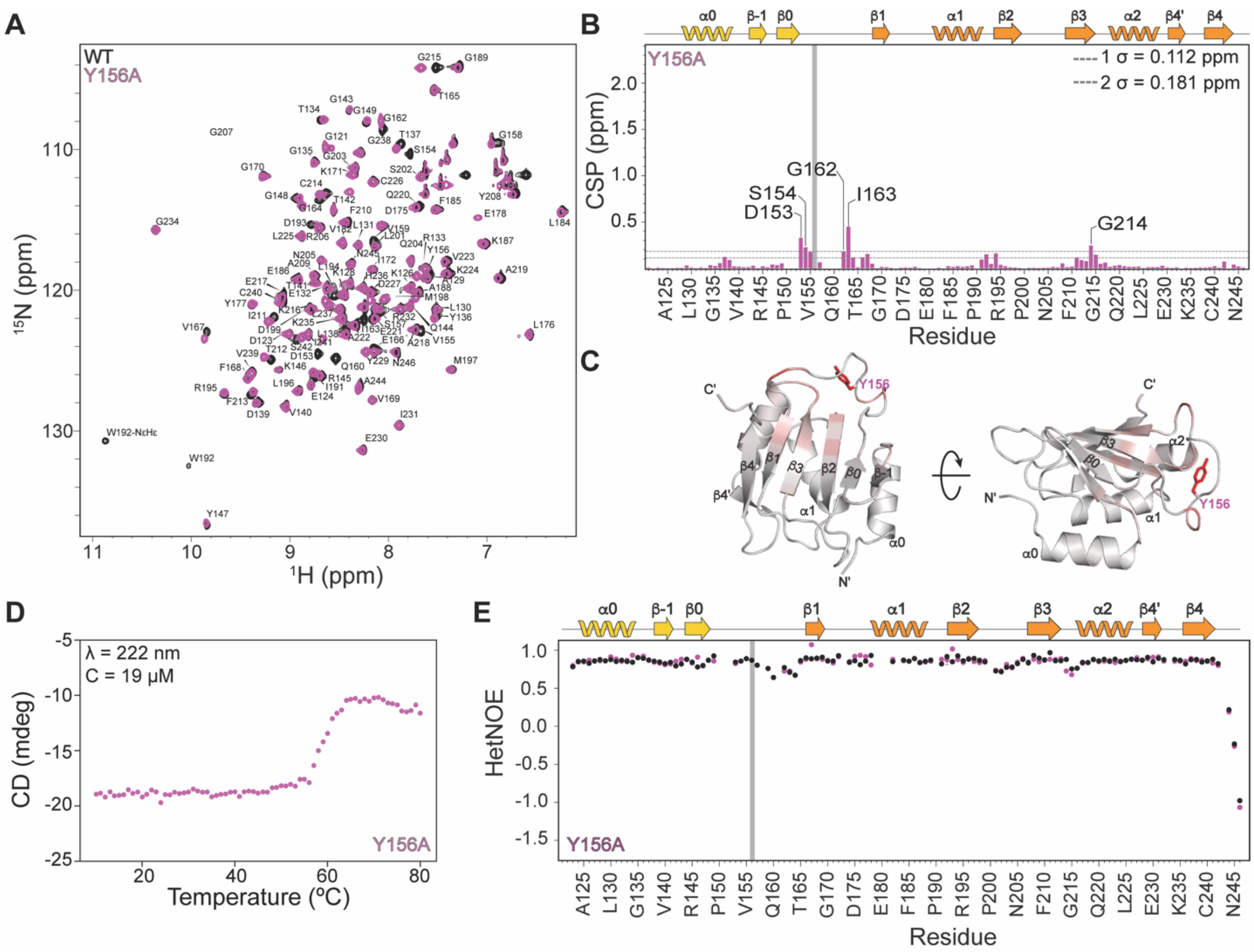


**Figure S12.** NMR and CD analysis of Y156A structure and thermal stability. A) Overlay of $^{1}$H-$^{15}$N HSQC spectra comparing backbone amide resonances of Y156A and WT; B) Y156A CSP plot; C) Y156A CSPs mapped onto the X-ray crystal structure; D) Representative raw CD melting trace of Y156A; E) Overlay of $^{1}$H-$^{15}$N heteronuclear NOE plot of Y156A (colored pink) and WT (colored black). The substitution site is shown as a gray bar.

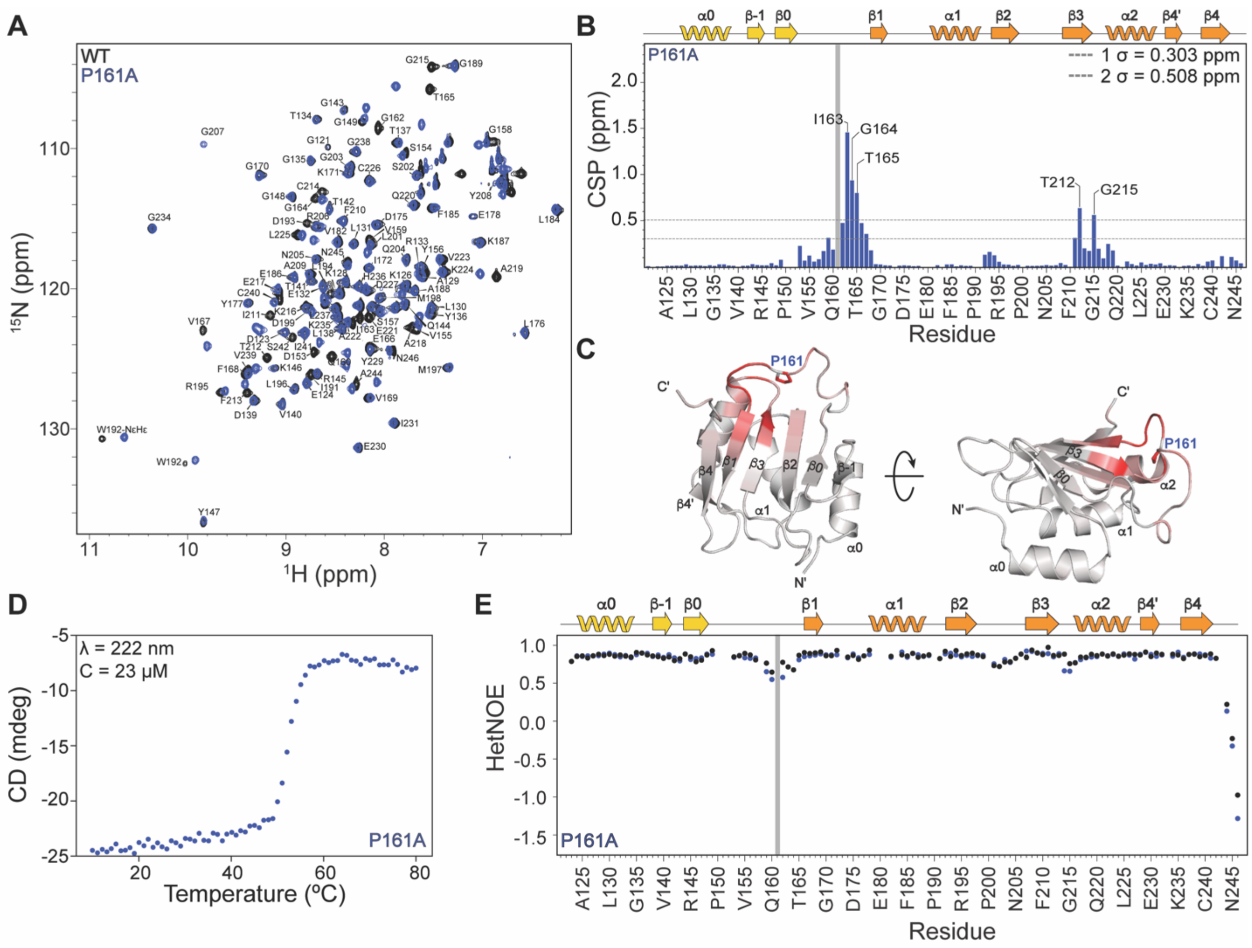


**Figure S13.** NMR and CD analysis of P161A structure and thermal stability. A) Overlay of $^{1}$H-$^{15}$N HSQC spectra comparing backbone amide resonances of P161A and WT; B) P161A CSP plot; C) CSPs mapped onto the X-ray crystal structure; D) Representative raw CD melting trace of P161A; E) Overlay of $^{1}$H-$^{15}$N heteronuclear NOE plot of P161A (colored marine) and WT (colored black). The substitution site is shown as a gray bar.

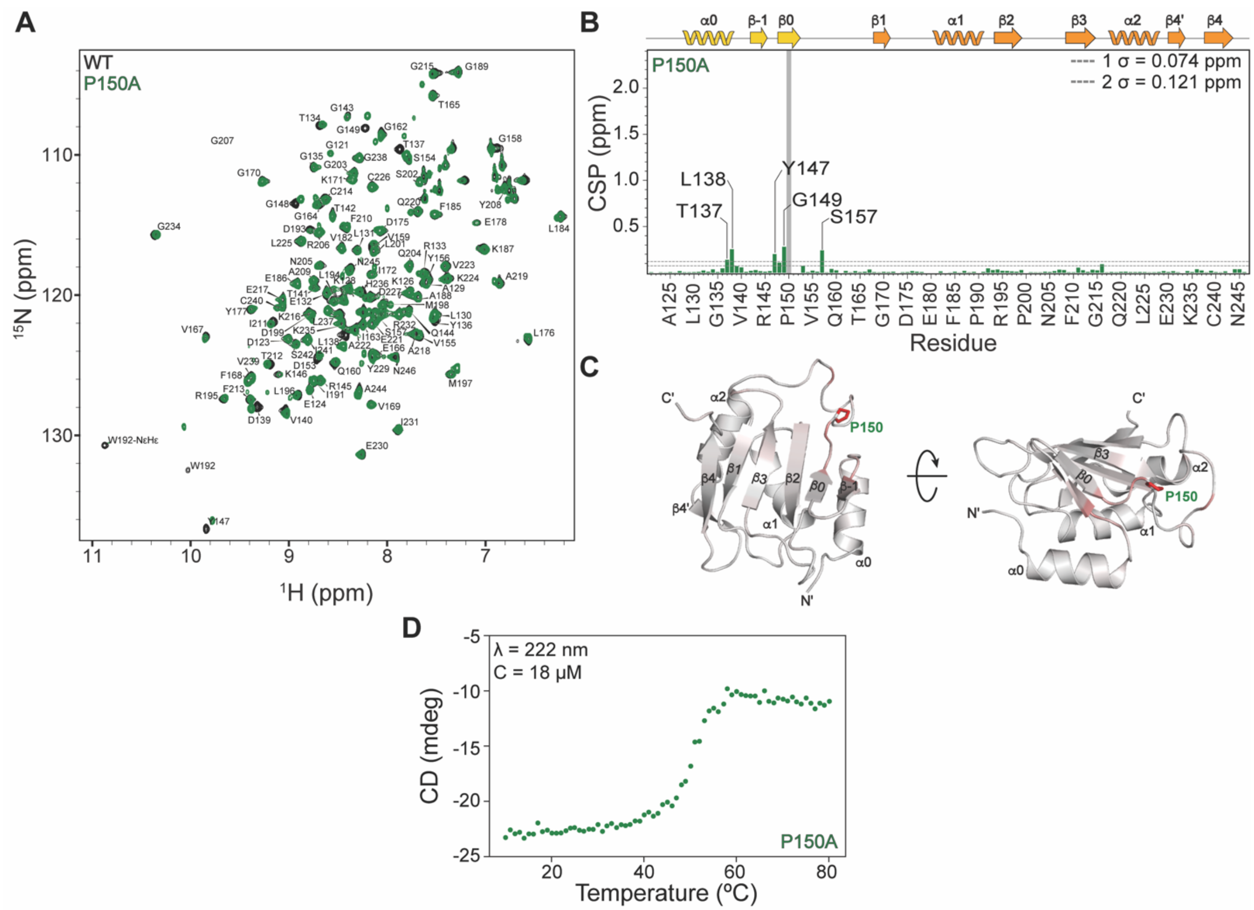


**Figure S14.** NMR and CD analysis of P150A structure and thermal stability. A) Overlay of $^{1}$H-$^{15}$N HSQC spectra comparing backbone amide resonances of P150A and WT; B) P150A CSP plot; C) CSPs mapped onto the X-ray crystal structure; D) Representative raw CD melting trace of P150A. The substitution site is shown as a gray bar.

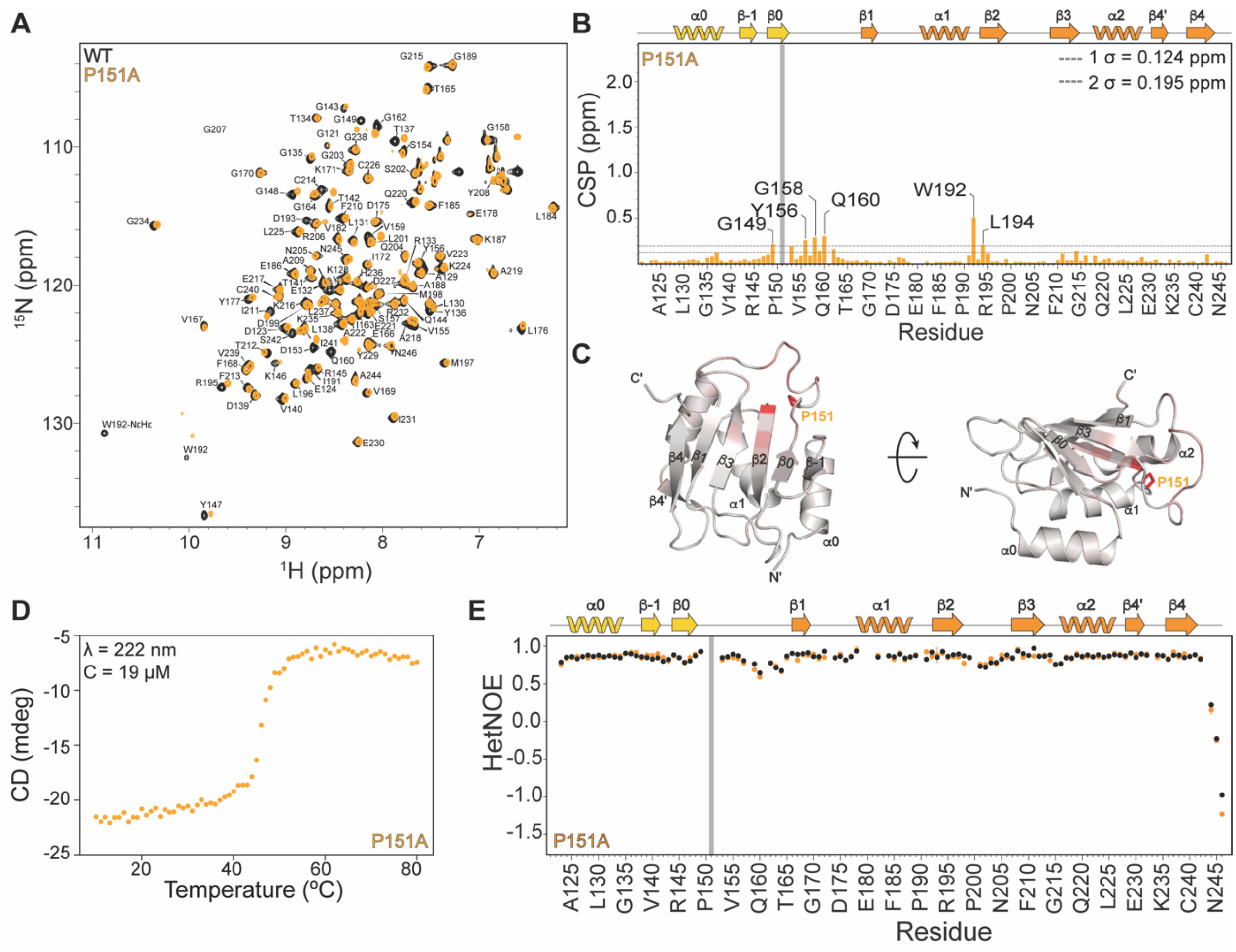


**Figure S15.** NMR and CD analysis of P151A structure and thermal stability. A) Overlay of $^{1}$H-$^{15}$N HSQC spectra comparing backbone amide resonances of P151A and WT; B) P151A CSP plot; C) CSPs mapped onto the X-ray crystal structure; D) Representative raw CD melting trace of P151A; E) Overlay of $^{1}$H-$^{15}$N heteronuclear NOE plot of P161A (colored orange) and WT (colored black). The substitution site is shown as a gray bar.

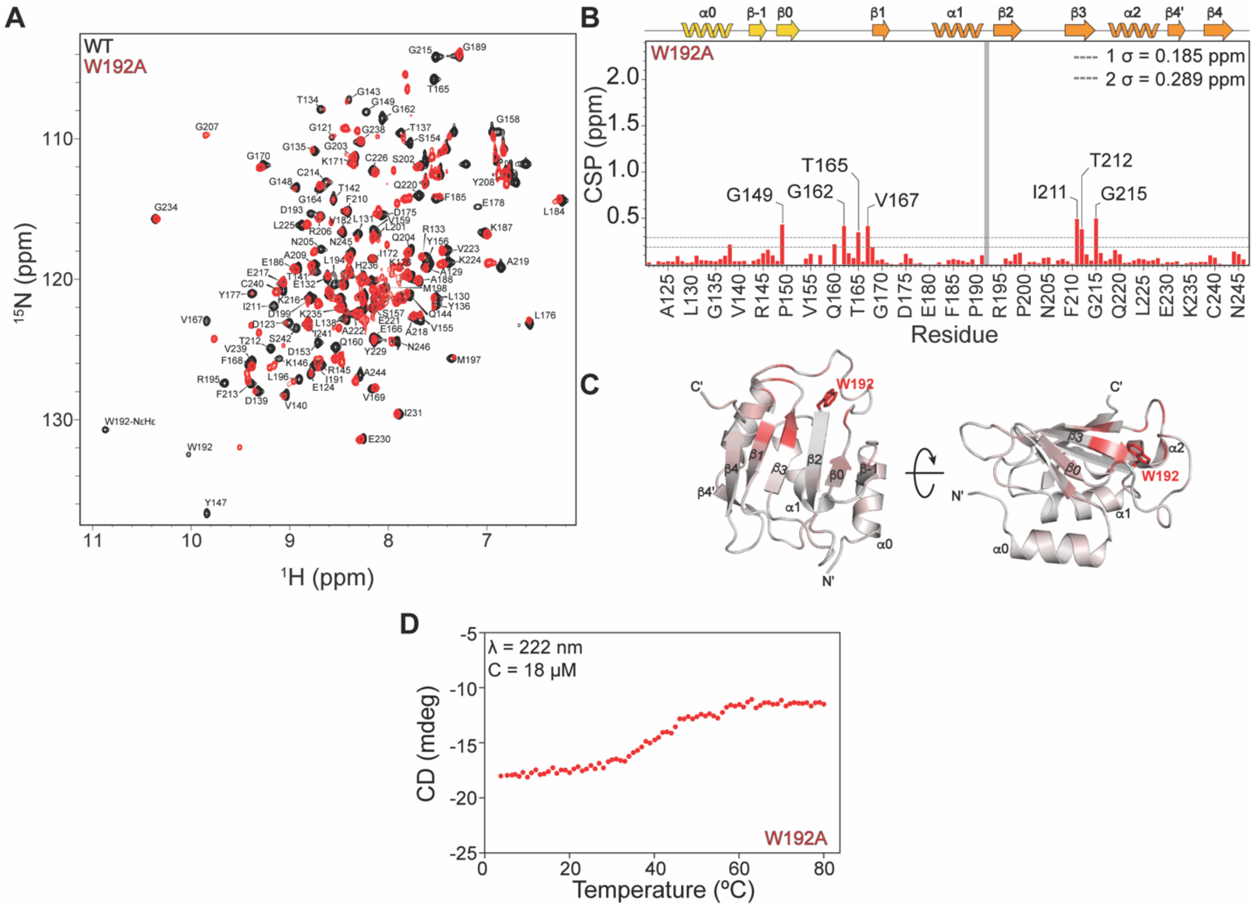


**Figure S16.** NMR and CD analysis of W192A structure and thermal stability. A) Overlay of $^{1}H$-$^{15}N$ HSQC spectra comparing backbone amide resonances of W192A and WT; B) W192A CSP plot; C) CSPs mapped onto the X-ray crystal structure; D) Representative raw CD melting trace of W192A. The substitution site is shown as a gray bar.

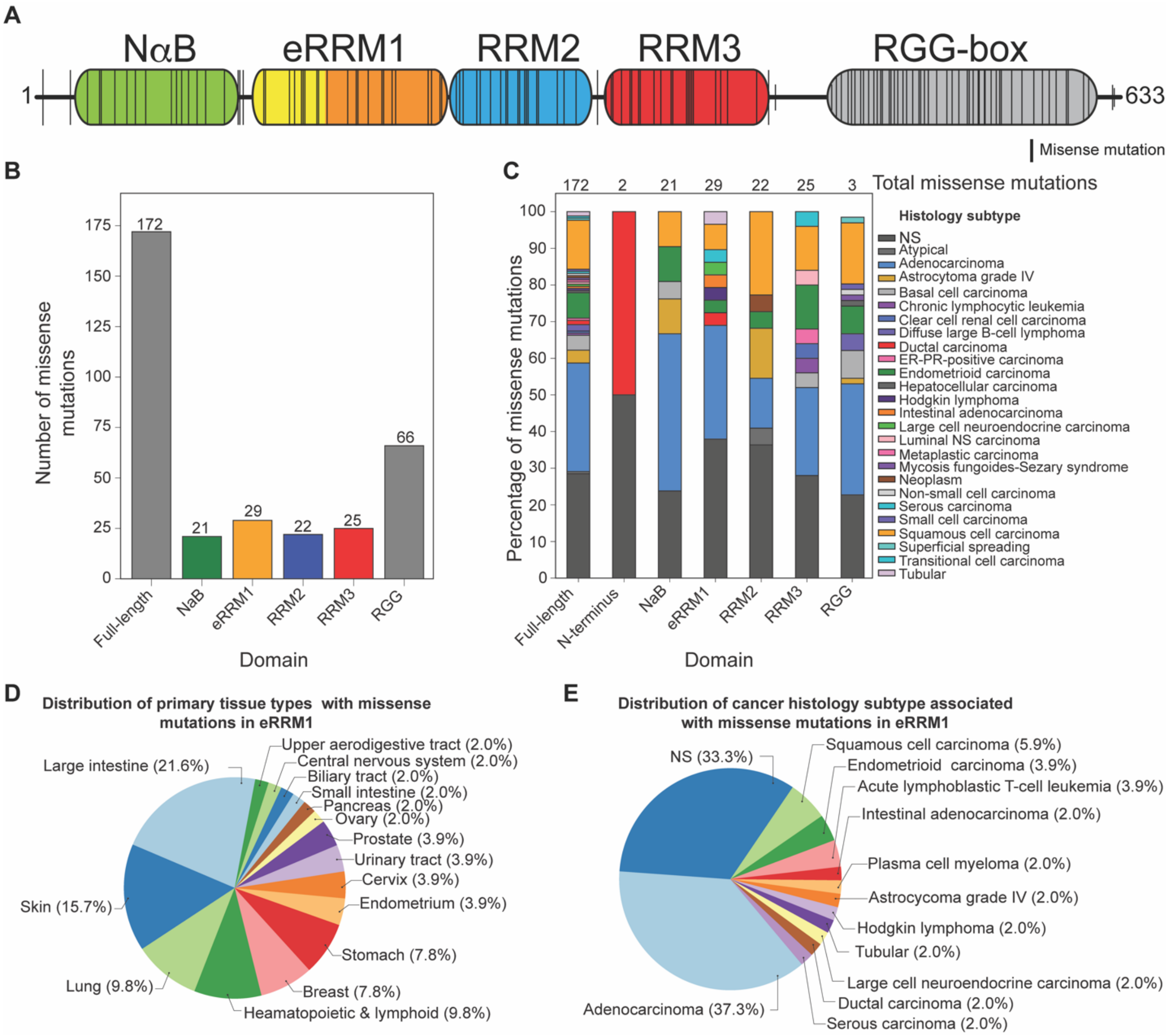


**Figure S17.** Cancer-associated somatic mutations in hnRNPR. A) Missense mutations from survey of COSMIC database, mapped onto the full-length hnRNPR domain topology; B) Distribution of missense mutations in hnRNPR domains with count shown above each bar; C) Distribution of cancer histology subtypes in hnRNPR domains with count shown above each bar; D) Distribution of primary tissue types with missense mutations in eRRM1; E) Distribution of cancer histology subtypes associated in eRRM1.

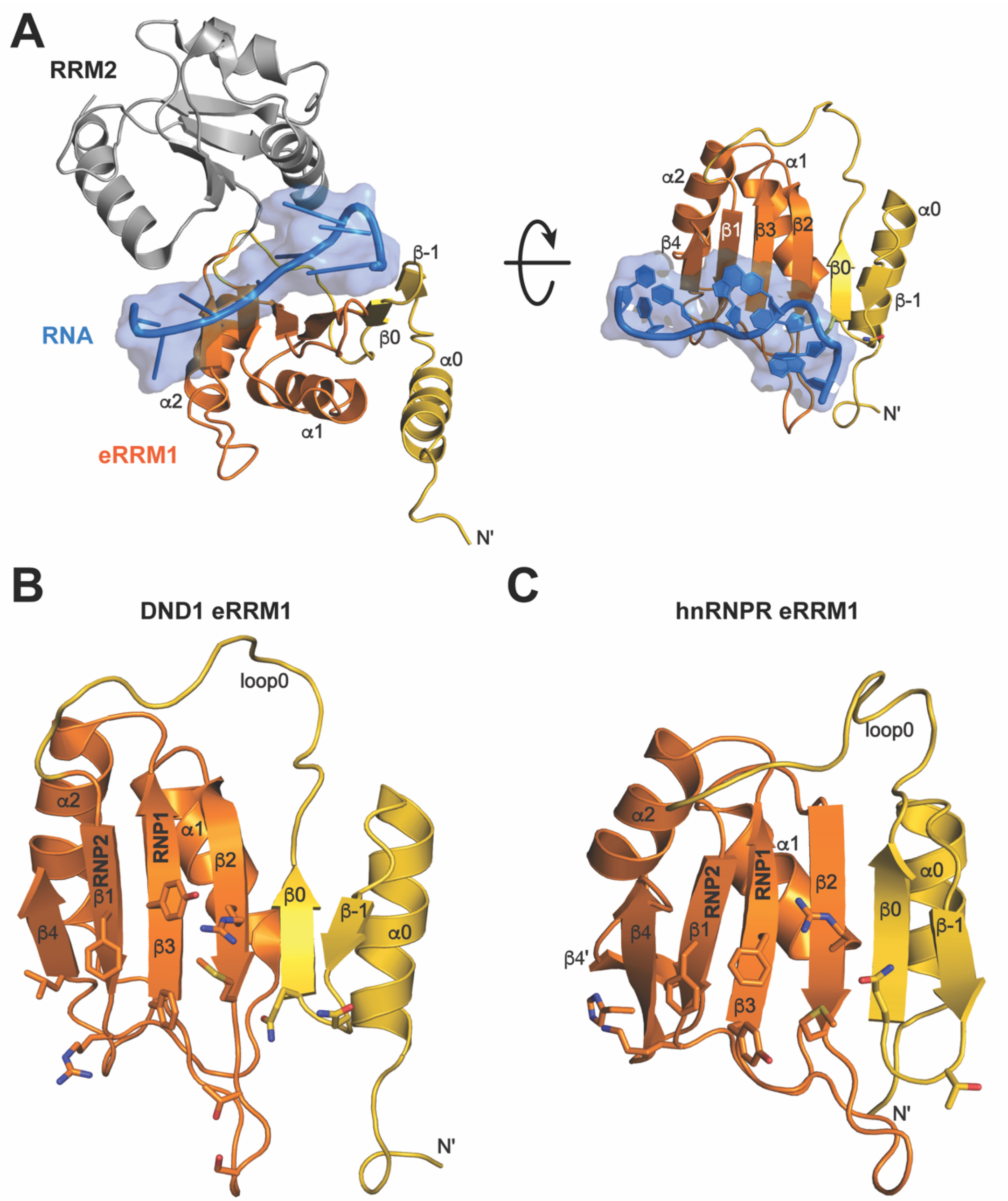


**Figure S18.** RNA recognition of DND1 eRRM1. A) DND1 RNA recognition shows the involvement of $N_{ext}$ β-hairpin in RNA recognition (PDB ID 7Q4L). RNA substrate is colored blue with surface and cartoon representation, eRRM1 is colored gold ($N_{ext}$) and orange (RRM), and RRM2 is colored gray. B) DND1 eRRM1 residues involved in RNA recognition, which include β3 (RNP1), β1 (RNP2), and residues on β2 and the $N_{ext}$ β-hairpin, are shown in stick representation. C) DND1 eRRM1 panel B residues at equivalent positions in hnRNPR eRRM1 are shown in stick representation.
